# Functional Synaptic Interactions and Inhibitory Circuitry of the PreBötzinger Complex in the Rhythmic Slice

**DOI:** 10.64898/2026.05.23.727419

**Authors:** Yaroslav I. Molkov, Hidehiko Koizumi, Jeffrey C. Smith

## Abstract

The preBötzinger Complex (preBötC) within the medulla oblongata contains neuronal circuits critical for generating the mammalian respiratory rhythm, but the functional connectivity among its core excitatory and inhibitory populations remains debated. Defining this connectivity requires disentangling synaptic interactions of functionally identified excitatory and inhibitory preBötC neurons with various electrophysiological phenotypes. We applied a novel synaptic conductance inference method to whole-cell recordings from genetically specified VgluT2-expressing (excitatory) and VGAT-expressing (inhibitory) preBötC neurons active in the rhythmic medullary slice in vitro, which contains core inhibitory-excitatory circuitry with an excitatory rhythmogenic kernel. We found that this circuitry consists of a self-exciting inspiratory VgluT2 population coupled to inspiratory and expiratory VGAT populations that interact reciprocally through inhibition. The functional inhibitory connectome is more complex than previously understood. However, compared with functional synaptic interactions inferred from recordings in the preBötC in situ, the neuronal synaptic conductance profiles in the rhythmic slice reveal a functionally reduced inhibitory connectome, characterized by prominent tonic expiratory inhibition and phasic inspiratory inhibition, without the characteristic multiphasic structure in situ. These results indicate that the functional excitatory and inhibitory circuit interactions within the preBötC isolated in vitro, although reduced relative to more intact states in situ, are intrinsically designed to generate coordinated inspiratory and expiratory population activity. Tonic expiratory phase inhibition together with inspiratory phasic inhibition serves to regulate excitability and phase transitions of the excitatory rhythmogenic kernel.

## Introduction

The preBötzinger Complex (preBötC) in the medulla oblongata is the primary site required for respiratory rhythm generation in mammals (Smith et al., 1991). This medullary region contains microcircuits with a heterogeneous collection of excitatory and inhibitory neuronal populations that exhibit inspiratory and expiratory patterns of neuronal activity (e.g., Schwarzacher et al., 1995; Ausborn et al., 2018). These circuits exhibit autorhythmic properties and generate rhythmic activity when isolated in rodent living medullary slices in vitro, providing a powerful experimental system to investigate the intrinsic rhythmogenic cellular and circuit mechanisms involved (e.g., Butera et al., 1999a,b; Smith et al., 2000; Feldman and Del Negro, 2006). While structural-functional properties of the resident excitatory and inhibitory populations have been analyzed to some extent (Koizumi et al., 2013), including circuit architectural features and intrinsic electrophysiological properties, defining the circuit functional organization—specifically, how excitatory and inhibitory populations synaptically interact to generate robust rhythmicity—remains a fundamental problem. A major hurdle has been the difficulty of disentangling excitatory and inhibitory synaptic drives during the respiratory cycle in phenotypically defined excitatory and inhibitory neuron types (Koizumi et al., 2013). Furthermore, how these synaptic drives in isolated circuits in vitro relate to preBötC circuit operation in more intact states of the brainstem respiratory network has not been directly addressed in experimental studies.

Most experimental slice studies have focused on burst patterns rather than quantifying the underlying synaptic conductances. Consequently, prior computational models have often invoked specific inhibitory or excitatory mechanisms—such as tonic inhibition or phase-specific synaptic interactions—without direct conductance-level experimental verification. The present study addresses this gap by providing conductance-resolved functional mapping of synaptic interactions and explicitly quantifying how synaptic interactions differ between the slice and intact network states.

In a recent study, we developed a novel computational inference technique (Molkov et al., 2025) to decompose total synaptic conductance into its excitatory and inhibitory components in single neurons using neuronal current-clamp recordings. By applying this method to neurons in the in situ arterially perfused mature rodent brainstem-spinal cord preparation with active respiratory circuits, we successfully mapped the dynamic “synaptic fingerprints” of respiratory neurons. This neuronal conductance profiling reveals the functional connectome between excitatory and inhibitory populations that underpins the dynamic operation of the in situ respiratory network.

However, the in situ preparation preserves a high degree of network complexity and external inputs. The acute medullary slice preparation remains the gold standard for investigating the minimal, intrinsic circuitry necessary for rhythm and inspiratory pattern generation. It is widely hypothesized that the slice contains a “reduced” circuit—a kernel of the network that is capable of sustaining rhythmicity in the absence of wider brainstem and feedback loops (e.g., Smith et al., 2000; Feldman and Del Negro, 2006; Smith et al., 2007, 2013; Del Negro et al., 2018). Yet, defining the functional connectivity of this reduced circuit, and how it differs from the intact network, requires rigorous quantitative analysis of synaptic interactions.

In this study, we apply our conductance inference methodology to the rhythmic slice preparation to define the core functional organization of the preBötC circuits. We obtained whole-cell recordings from genetically identified preBötC VgluT2-expressing glutamatergic and VGAT-expressing inhibitory neurons in slices from transgenic mice, combined with computational analysis to reconstruct the dynamic synaptic inputs of these neurons.

Our analysis reveals a distinct, reduced circuit configuration active in the preBötC within the slice. We identify four functionally active populations based on neurotransmitter identity and neuronal firing phase. Crucially, we demonstrate that while inspiratory rhythm is driven by excitatory interactions within the synaptically coupled VgluT2+ population, the phasic transitions are enforced by specific reciprocal inhibitory interactions between VGAT+ populations. Furthermore, unlike in the intact in situ rat brainstem-spinal cord preparation, the expiratory populations active in the slice lack the synaptic inputs and interactions required to produce multiphasic expiratory patterns. These findings allow us to define a reduced functional state of the preBötC circuits, in which a self-exciting inspiratory source is coupled with a reciprocal inhibitory motif, regulated by tonic expiratory phase inhibition, that constitutes the basic circuit interactions for respiratory rhythmic pattern generation and control in vitro.

## Methods

### Animals, slice preparation, and cell targeting

All animal procedures were approved by the Animal Care and Use Committee of the National Institute of Neurological Disorders and Stroke (Animal Study Proposal 1154–21).

Experiments were performed in rhythmically active medullary slice preparations from neonatal (postnatal day 3–8) transgenic mice of either sex, as described previously for studies targeting genetically identified preBötC excitatory or inhibitory neurons validated in reduced respiratory preparations (Koizumi et al., 2013; Ausborn et al., 2018). Excitatory neurons were targeted for whole-cell recording in Slc17a6-Cre;Rosa-tdTomato mice (referred to here as VgluT2-tdTomato mice), and inhibitory neurons were targeted in VGAT-Cre;Rosa-tdTomato mice (VGAT-tdTomato mice). TdTomato-positive preBötC neurons in the slice were identified by optical live fluorescence imaging.

Rhythmically active slices (300–400 *µ*m) were superfused at 4 ml/min with artificial cerebrospinal fluid containing (in mM): 124 NaCl, 25 NaHCO_3_, 3 KCl, 1.5 CaCl_2_, 1.0 MgSO_4_, 0.5 NaH_2_PO_4_, and 30 D-glucose, equilibrated with 95% O_2_/5% CO_2_ (pH 7.35–7.40, 27*^◦^*C). Rhythmic respiratory activity was maintained by elevating extracellular K^+^ to 8–9 mM. Inspiratory motor output was monitored from hypoglossal (XII) rootlets with suction electrodes, amplified 50,000–100,000×, band-pass filtered at 0.3–2 kHz, and digitized at 10 kHz.

Due to variability in the amplitude and stability of the XII motor output across slices, we treated the XII recording primarily as a cycle-timing reference and applied additional preprocessing safeguards (below) to ensure reliable phase detection.

Optical imaging was performed with a Leica TCS SP5 II MP upright microscope equipped with a 20× water-immersion objective (NA 1.0) and a Ti:Sapphire laser (800–880 nm) for two-photon fluorescence imaging. Whole-cell current- and voltage-clamp recordings were obtained from the imaged excitatory or inhibitory neurons with a HEKA EPC-9 amplifier using borosilicate electrodes (4–6 MΩ) filled with (in mM): 130 K-gluconate, 5 Na-gluconate, 3 NaCl, 10 HEPES, 4 Mg-ATP, 0.3 Na-GTP, and 4 sodium phosphocreatine, adjusted to pH 7.3 with KOH. Measured voltages were corrected for a –10mV liquid-junction potential. Series resistance was compensated online by approximately 80% and adjusted during the recording. Voltage-clamp recordings with obvious space-clamp failure were excluded before quantitative analysis.

### Recording inventory, acquisition modes, and acceptance screening

The data repository (see Code and Data Availability) contains 59 analyzed current- and voltage-clamp recordings spanning four phenotype groups: VgluT2 inspiratory (VgluT2-I), VgluT2 expiratory (VgluT2-E), VGAT inspiratory (VGAT-I), and VGAT expiratory (VGAT-E). In most cases, the recording data files for each neuron contain both current-clamp (C) and voltage-clamp (V) recordings, whereas some files contain only current- or voltage-clamp recordings when, for technical reasons, both recording modes were not obtained for that neuron. Raw neuronal and XII electrophysiological signals were originally digitized at 10 kHz and then downsampled to 1 kHz for the exported analysis files used in this repository. This downsampling preserved the slower respiratory-cycle modulation, the stepwise command epochs, and the phase-binned current–voltage relationships used for conductance inference; reconstruction of the detailed spike waveform was not required for the analysis. Each exported trace contained four aligned columns: time, current channel, voltage channel, and integrated XII activity. In current clamp, the current and voltage channels corresponded to the injected current and membrane potential, respectively; in voltage clamp, they corresponded to the measured holding current and the command voltage.

Because of the variability of the number of current/voltage steps, total duration, and stability of the recordings across slices, to select recording epochs for analysis, we did not impose rigid file-level requirements, such as at least three usable command steps or a prescribed current/voltage span, as in the companion in situ study of Molkov et al. (2025). Instead, we used an inclusive recording-level screen and then applied robust outlier filtering only after conductance reconstruction. The only strict acceptance criterion imposed for inclusion in the population summaries was that the analyzed segment contain more than 25 detected XII cycles across the retained interval. This yielded 50 accepted recordings (27 current clamp, 23 voltage clamp): 20 VgluT2-I, 12 VGAT-I, 17 VGAT-E, and 1 VgluT2-E recording (Table 1) from 31 neurons total. Of these 31 neurons, 18 contributed paired current- and voltage-clamp recordings of the same cell to the accepted set, 11 contributed a single recording mode, 1 contributed two same-mode recordings, and 1 yielded only sub-threshold recordings and was excluded entirely by the cycle-count screen. The *N* values in summary plots and tables therefore refer to the total number of recordings (C and V) for each phenotype group, not to individual cells. We treated C- and V-clamp files as separate analysis units because recording-mode-to-mode variability was comparable to cell-to-cell variability, and the two modes yielded qualitatively consistent conductance estimates (see Figure 4 below).

**Table 1:**
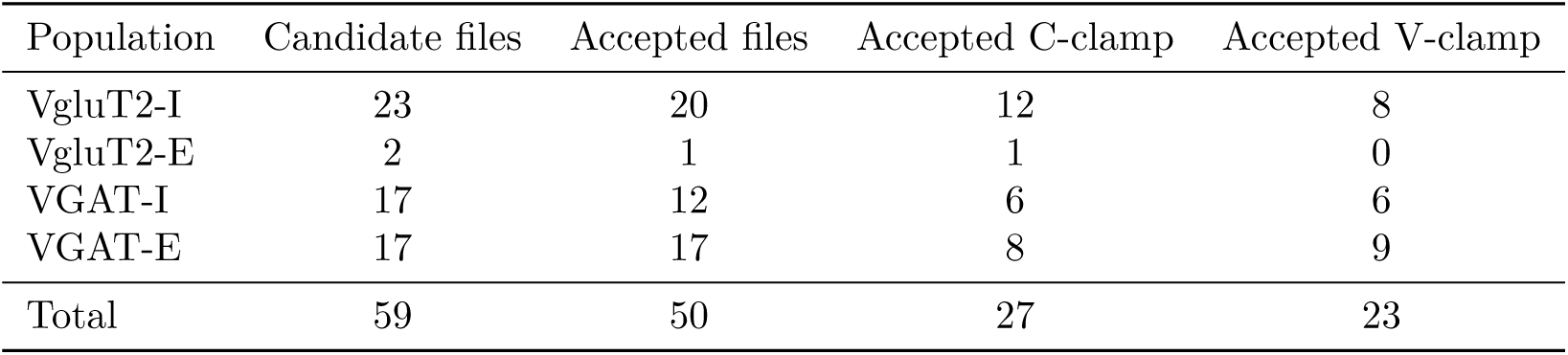
Recording inventory in the repository and after automated cycle-count screening. . Counts refer to recordings, not unique neurons.

The command protocols used for conductance inference consisted of piecewise-constant current steps in current-clamp and piecewise-constant holding-voltage steps in voltage-clamp (Figure 1). A representative current-clamp/voltage-clamp pair from the same VgluT2-I neuron is shown to make the stepped acquisition strategy explicit. Across the selected recording interval (Figure 1, blue regions), the multiple command levels were not analyzed one step at a time; instead, samples from all usable steps were pooled within each respiratory-phase bin, so that the full recording collectively provided the current–voltage spread needed for

**Figure 1:**
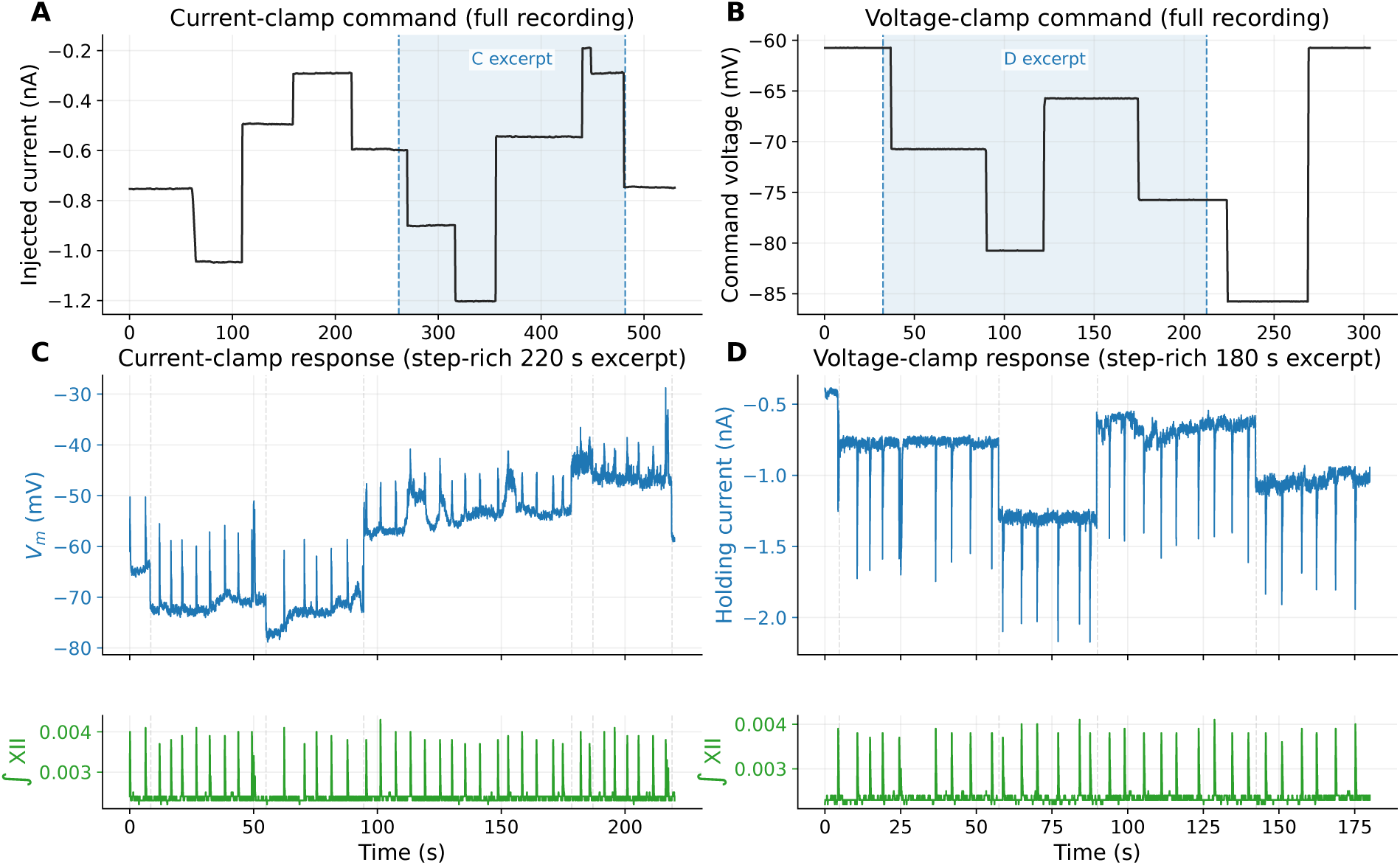
**Representative stepped acquisition protocols used for conductance inference**. **A**, full current-clamp command trace for VgluT2-I-Cell2-C, shown after coarse median smoothing, to highlight the piecewise-constant current injections; the blue shaded interval marks the recording segment expanded in panel **C**. **B**, full voltage-clamp command trace for the matched recording VgluT2-I-Cell2-V; the blue shaded interval marks the segment expanded in panel **D**. **C–D**, expanded recording excerpts selected automatically for analyses; the response channel is plotted in the upper lane (membrane potential in current clamp, holding current in voltage clamp) and the integrated XII reference is plotted separately in the lower lane. Gray dashed lines mark command transitions that fall inside the excerpt. The full step-command traces in **A** and **B** were used to sample a range of current–voltage operating points across many respiratory cycles.

### Manual curation and preprocessing of analyzed recording segments

Several recordings required limited manual curation before automated inference of conductances. The purpose of this step was to isolate the longest interval that was reasonably stationary in both the electrophysiological channels and the XII respiratory cycle reference, rather than to select epochs with any particular conductance pattern. Files were therefore trimmed at the beginning, when the recording was still stabilizing; at the end, when rhythmicity or seal quality deteriorated; and when instability was present at both ends, to the middle portion that remained acceptably stationary. Thus, “middle selection” was implemented by defining start and end boundaries around a single contiguous interval rather than by stitching together disjoint fragments. In the current repository version, 6 accepted recordings used a non-zero starting offset, and 6 used an explicit end truncation. These edits were intentionally conservative and were applied before any conductance estimates were calculated.

All signal processing was then automated. The current and voltage channels were median-filtered with a per-file half-width defined in the analysis configuration (default 25 samples, corresponding to a 51-sample window). Because the mouse XII rootlet signal was not always sufficiently robust on its own, the reference channel was optionally combined with the simultaneously recorded electrophysiological channel when the XII trace was too weak, noisy, or unstable to give reliable cycle onsets. In those cases, the reference channel was converted to a local z-score and linearly combined with a local z-score of the membrane-potential channel (current clamp) or the current channel (voltage clamp): *p*_mix_(*t*) = *z*[*p*(*t*)] + *sz*[*y*(*t*)], where *p*(*t*) is the integrated XII signal, *y*(*t*) is the auxiliary electrophysiological channel, and *s* = ±1 was chosen empirically to align the sign of the auxiliary signal with the inspiratory onset. Local means and standard deviations were computed in a sliding window (typically 2000 samples). This mixed signal was used only for cycle detection and phase assignment; conductance inference itself still used the original current and voltage channels. Among the 50 accepted recordings, 6 used this mixed trigger signal, and 12 used a broader median post-filter on the final reference trace than the default setting.

### Cycle detection and phase normalization

Accurate determination and normalization of the respiratory cycle phase are critical for our cycle-phase-based conductance analyses. The cycle phase was derived from the processed XII reference trace. Let *p*(*t*) denote the final reference signal after optional mixing and median filtering. The detection threshold was chosen automatically from the median and robust peak amplitude of the full trace:

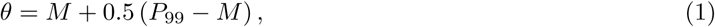

where *M* = median(*p*) and *P*_99_ is the 99th percentile of *p*. A cycle onset *T_n_* was defined as a rising threshold crossing, *p*[*i* − 1] *< θ* ≤ *p*[*i*], that remained above threshold for at least 10 consecutive samples. Crossing time was linearly interpolated between the two bracketing samples. Successive onsets were required to be at least 2 s apart to avoid double-counting within the same inspiratory burst. Phase was then normalized linearly between successive accepted onsets:

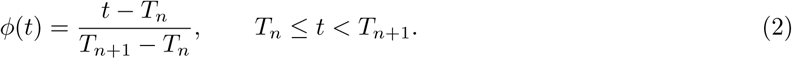

Thus, *ϕ* = 0 corresponds to the XII burst onset and *ϕ* ∈ [0, 1) indexes the respiratory cycle independently of its absolute duration.

### Phase-binned current–voltage regression

Conductance inference followed the phase-domain method introduced in the companion in situ study of Molkov et al. (2025). We write the membrane-current balance as

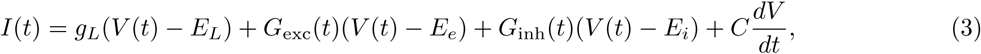

where *I*(*t*) is the current channel, *V* (*t*) is the voltage channel, *g_L_* and *E_L_* are the effective leak conductance and reversal potential, *G*_exc_ and *G*_inh_ are the phase-dependent excitatory and inhibitory conductances, and *E_e_* and *E_i_* are their reversal potentials. In this formulation, *g_L_* and *E_L_* absorb both intrinsic leak and any approximately tonic background synaptic drive, allowing the dynamic respiratory modulation to be isolated in *G*_exc_(*ϕ*) and *G*_inh_(*ϕ*).

Assuming that synaptic conductances vary primarily with respiratory phase and that the capacitive term is small relative to the synaptic terms on this time scale, Equation 3 reduces within each phase bin to a linear current–voltage relation,

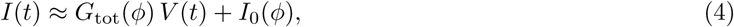

where *G*_tot_(*ϕ*) = *g_L_* + *G*_exc_(*ϕ*) + *G*_inh_(*ϕ*) and *I*_0_(*ϕ*) is the phase-specific intercept.

The normalized cycle was discretized into 1000 phase bins. For each bin, all samples whose phases fell inside that bin were pooled across the analyzed segment. If more than two points were available, an initial linear regression was fitted and residuals were computed. Points with residual magnitude greater than 2.5 standard deviations were excluded, and the regression was recomputed on the retained points. Bins with fewer than three retained points were discarded.

The implementation differed slightly between acquisition modes to improve numerical stability while preserving the same *I*–*V* output. In voltage clamp, the final regression was fitted directly as *I* = *aV* + *b*, so the slope and intercept were simply *G*_tot_ = *a* and *I*_0_ = *b*. In current clamp, the second-pass fit was performed as *V* = *αI* + *β*, then algebraically transformed back to

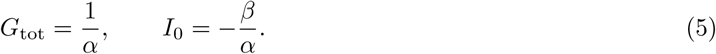

This avoids fitting a nearly vertical *I*–*V* relation in recordings where the injected current is the more naturally discretized variable.

### Leak estimation and decomposition into excitation and inhibition

For each valid phase bin, the fitted current–voltage relation defines the total conductance and intercept trajectory shown in Figure 2. The current expected at the inhibitory and excitatory reversal potentials is

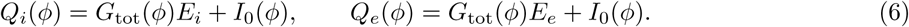

**Figure 2:**
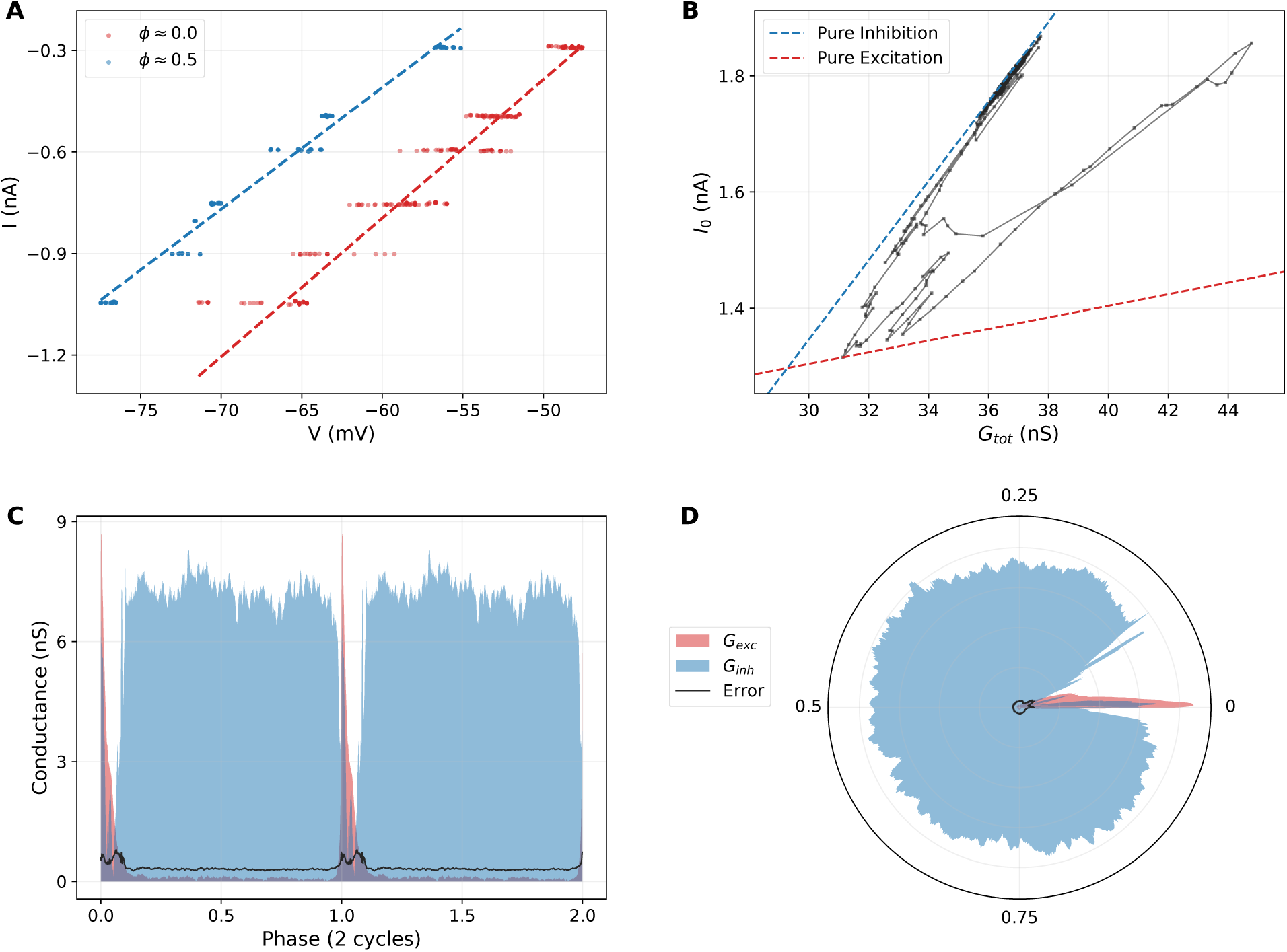
**Geometric conductance inference workflow**. **A**, representative phase-binned *I*–*V* regressions for two phases of the respiratory cycle (VgluT2-I-Cell2-C recording). The point clouds are organized into horizontal strips because each strip corresponds to repeated samples collected at the same injected-current step; the horizontal extent of a strip shows cycle-to-cycle variability in membrane voltage at that current level and respiratory phase. This within-step voltage variability contributes to scatter around the fitted *I*–*V* relation and therefore to the regression uncertainty propagated as the black error trace in **C** and **D**. **B**, wedge plot formed by the trajectory of (*G*_tot_*, I*_0_) across the cycle, bounded by the theoretical pure-excitation and pure-inhibition lines. **C**, reconstructed excitatory (red) and inhibitory (blue) conductances across two normalized cycles. **D**, the same conductances shown in polar coordinates over one normalized cycle.

Assuming that each recording contains at least one phase at which excitation is minimal and one phase at which inhibition is minimal, the leak-only currents at the inhibitory and excitatory reversal potentials are estimated as

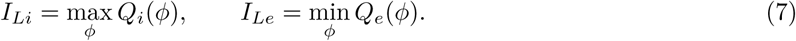

These values give the effective leak parameters

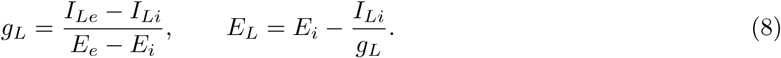

Excitatory and inhibitory conductances are then recovered as

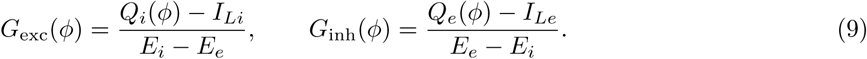

For reporting and plotting, conductances were normalized by the effective leak conductance *g_L_*, so all population summaries are expressed as *G*_exc_*/g*_leak_ and *G*_inh_*/g*_leak_.

### Data-driven estimation of inhibitory reversal potential

The conductance decomposition depends critically on the inhibitory reversal potential, so treating *E_i_*as a single fixed constant would force every recording to obey the same inhibitory geometry. Instead, we estimated *E_i_* separately for each recording from the wedge plot formed by the phase-binned regression outputs (*G*_tot_(*ϕ*)*, I*_0_(*ϕ*)).

Figure 2B illustrates the logic of this estimate. Each point on the black trajectory corresponds to one respiratory phase bin in the (*G*_tot_*, I*_0_) plane. The blue dashed line is the pure-inhibition boundary, and the red dashed line is the pure-excitation boundary. If a phase contains inhibition with negligible excitation, its point lies on the blue boundary. Adding excitation moves the point into the interior of the wedge, away from that boundary, but cannot move it above the blue line because excitatory conductance is constrained to remain nonnegative. The upper envelope of the trajectory in Figure 2B therefore reflects inhibition alone.

Because the slope of that upper boundary is set by −*E_i_*, the wedge geometry provides a direct recording-specific estimate of inhibitory reversal potential before excitatory and inhibitory conductances are separated.

In practice, after the robust phase-binned *I*–*V* regressions were computed, the analyzer searched for the tightest line that remained a global upper bound of all valid wedge points and took its slope as the recording-specific value of −*E_i_*. The resulting *E_i_* was then used in the subsequent conductance decomposition for that recording. The analyzer was initialized with *E_e_* = −10 mV and an initial inhibitory guess of *E_i_* = −80 mV; if too few valid regression points were available to define a reliable boundary, the supplied inhibitory value was retained rather than forcing a geometric estimate. In the accepted data set, however, the automatic estimate was applied to all 50 included recordings. This recording-specific procedure prevented the decomposition from depending on a single assumed inhibitory reversal potential and ensured that each inferred wedge remained internally consistent with the geometry of its own regression cloud.

### Definition of inspiratory and expiratory summary metrics

The per-bin conductance traces were next reduced to scalar metrics for population analysis. First, the medians of *G*_inh_(*ϕ*) and *G*_exc_(*ϕ*) across all valid bins were computed, together with median absolute deviations scaled to a 3*σ*-equivalent threshold. Any phase bin whose inhibitory or excitatory conductance lay outside this robust threshold was marked as belonging to a transient interval. The longest consecutive transient chain on the circular phase axis was then identified as the phasic respiratory event, and the complementary bins were treated as the stationary interval.

For each recording, the analyzer exported (i) the mean inhibitory and excitatory conductances over the stationary interval and (ii) the conductances in the onset-aligned bin at the beginning of the XII-defined inspiratory burst. Because the inspiratory conductance profiles in these recordings were decrementing after burst onset, this onset-aligned value was used as a practical estimate of peak inspiratory conductance for the bar plots and summary tables, whereas the stationary-interval means were used as the expiratory/interburst metric.

### Population statistics and outlier handling

Population summaries were produced separately for VgluT2-I, VgluT2-E, VGAT-I, and VGAT-E recordings. After the cycle-count screen, one row per accepted recording was written to a group-specific CSV file containing the stationary means and onset-aligned peak-inspiratory conductances. Outlier handling was deliberately postponed to this stage rather than applied as a strict recording-level exclusion rule.

Outliers were handled by a standard robust two-stage procedure: calculate a provisional distribution from all accepted recordings, identify extreme points using quantile-based criteria, remove those points from the summary statistics, and then recalculate the reported mean and SEM from the retained inliers. In practice, for each group and each scalar metric, quartiles were computed across recordings, the interquartile range was defined as IQR = *Q*_3_ − *Q*_1_, and recordings outside the standard Tukey interval [*Q*_1_ − 1.5 IQR*, Q*_3_ + 1.5 IQR] were labeled as outliers. Mean and SEM were then recomputed from the retained inliers only. Those outliers are excluded from the summary bars. This is a standard accepted way to summarize small, heterogeneous experimental data sets while limiting the influence of rare extreme values. We adopted this two-stage strategy because the slice data set contains some recordings with limited step-command span or weaker reference signals that would be lost under rigid step-count criteria even though their reconstructed conductances fall well within the physiological distribution of the corresponding population.

### Software implementation

The conductance inference algorithm—including median filtering, network-cycle detection, phase binning, robust phase-binned *I*–*V* regression, automatic estimation of *E_i_*, and decomposition of total conductance into excitatory and inhibitory components—was implemented as a single C++11 program compiled with g++ -O3 and depending only on the C++ standard library. Batch processing across recordings, group-level statistical summaries, outlier screening, and the generation of all manuscript figures were implemented in Python 3 using NumPy, SciPy, and Matplotlib. The full pipeline from raw current and voltage traces to the figures and tables presented in this paper is reproducible from a single Makefile.

## Results

### Firing Phenotypes and Inferred Synaptic Conductances

We analyzed synaptic inputs received by the rhythmically active respiratory neurons to understand how these inputs shape their firing phenotypes (Figure 3). As illustrated, we found two main phenotypes: inspiratory (I) neurons spiking during XII motor output and expiratory (E) neurons spiking during the interval between inspiratory bursts and suppression of spiking activity during the inspiratory phase. Each spiking phenotype had subpopulations of excitatory (VgluT2-I, VgluT2-E) and inhibitory (VGAT-I, VGAT-E) neurons. While most of the I neurons began spiking synchronously with the onset of the integrated XII nerve discharge (e.g., VgluT2-I in Figure 3), some of the excitatory and inhibitory I neurons exhibited depolarization and spiking starting hundreds of milliseconds before (pre-I) the integrated XII nerve discharge (e.g., see VGAT-I neuron in Figure 3 and Supplemental Figure 3 for more examples). Matched voltage-clamp recordings confirm that this pre-I spiking is driven by a progressively developing net inward synaptic current. Furthermore, some of the E neurons exhibited pre-I membrane hyperpolarization (e.g., VGAT-E in Figure 3 and Supplemental Figure 4) and associated outward synaptic current.

**Figure 3:**
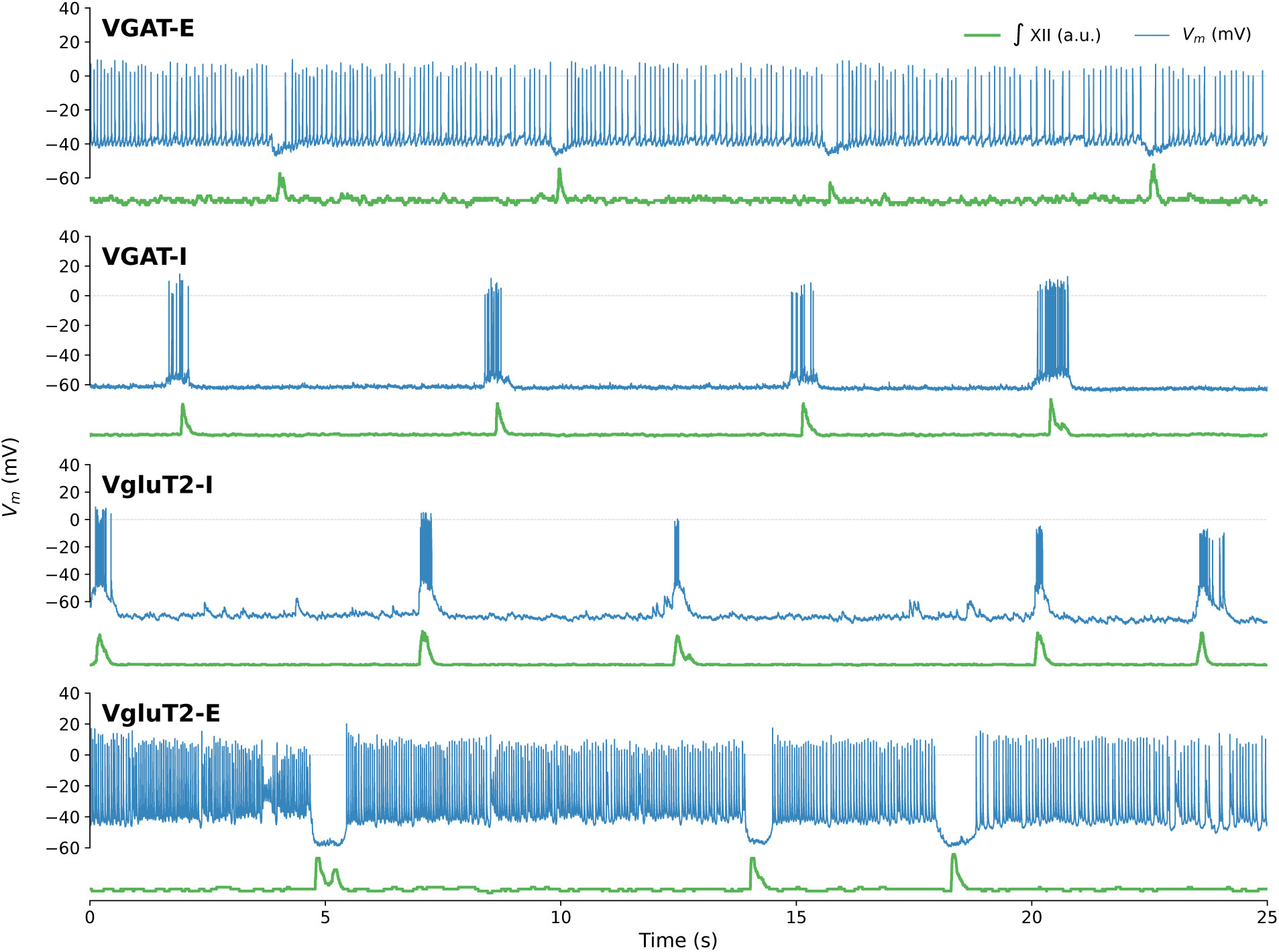
**Neuronal firing patterns**. Representative 25-second current-clamp recording episodes high-lighting the firing patterns of typical neurons in the VgluT2-I, VgluT2-E, VGAT-I, and VGAT-E populations. The blue trace illustrates the membrane potential, exhibiting distinct bursting dynamics that are time-aligned with the network rhythm. The green trace represents the integrated XII nerve activity (^∫^ XII), providing a global reference for the inspiratory phase. For the VgluT2-E example, the E-neuron spiking frequency reflects the level of injected depolarizing current; see Supplemental Figure 4 for additional examples of E-phase spiking activity.

**Figure 4:**
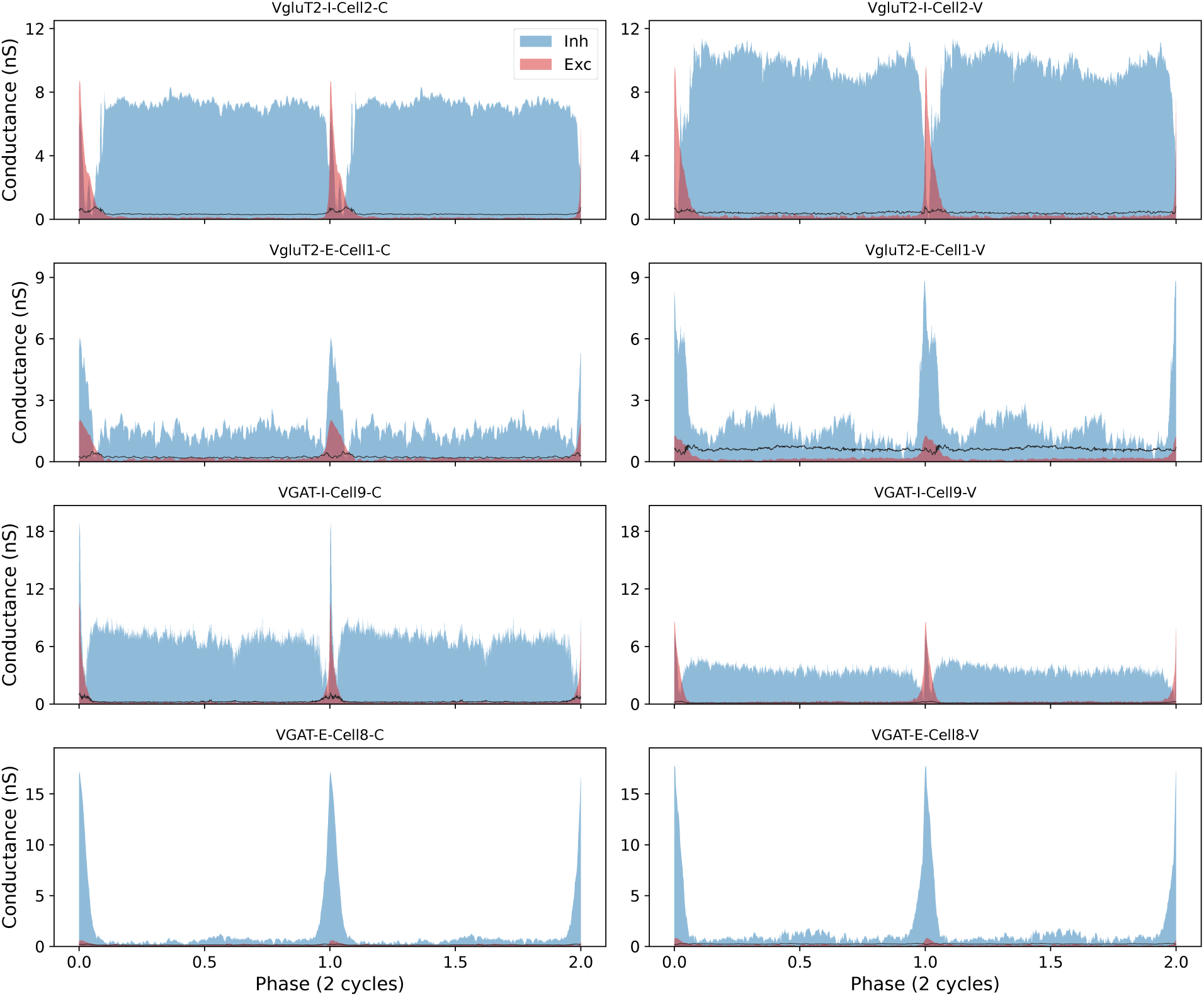
**Synaptic conductance profiles of representative excitatory and inhibitory neurons**. Absolute excitatory (red) and inhibitory (blue) synaptic conductances (in nS) estimated from current clamp (left column) and voltage clamp (right column) recordings. The traces show the phase-dependent mean conductance over two normalized respiratory cycles for four representative neurons: **A**. VgluT2+ inspiratory neuron (VgluT2-I-Cell2), **B**. VgluT2+ expiratory neuron (VgluT2-E-Cell1), **C**. VGAT+ inspiratory neuron (VGAT-I-Cell9), and **D**. VGAT+ expiratory neuron (VGAT-E-Cell8). Shaded areas indicate the mean conductance, and black lines indicate the standard error. Note the distinct temporal organization of synaptic inputs across inspiratory and expiratory cell types.

Figure 4 illustrates the inferred excitatory and inhibitory conductances for representative inspiratory and expiratory neurons, covering both excitatory (VgluT2+) and inhibitory (VGAT+) populations. It shows that synaptic conductance waveforms reconstructed from current-clamp and voltage-clamp recordings for each neuron type are largely consistent, indicating that the inference method yields comparable results across recording modes. In both configurations, the same phase-dependent structure of excitation and inhibition is preserved for each cell type: inspiratory neurons exhibit a transient peak of excitatory conductance during inspiration together with dominant tonic inhibitory conductance during expiration, whereas expiratory neurons receive strong phasic inhibition during inspiration and virtually no excitatory synaptic drive during their active phase. Absolute conductance magnitudes and temporal profiles are similar between recording modes, and the timing of conductance transitions aligns closely, supporting the validity of current-clamp– based conductance reconstruction. Minor quantitative differences in amplitude or variability likely reflect methodological factors, including imperfect space clamp and series resistance errors in voltage clamp, as well as deviations from the steady-state and linearity assumptions underlying the current-clamp inference (e.g., membrane capacitance effects, conductance nonstationarity, or phase jitter across cycles). Overall, the agreement between the results from the two recording methods indicates that the observed dominance of tonic expiratory inhibition and phase-specific excitation is a robust property of the network rather than an artifact of the recording configuration.

Inspiratory neurons consistently exhibited a specific pattern of synaptic input: they received phasic inspiratory excitation during the burst phase, prominent tonic expiratory inhibition during the inter-burst interval, with disinhibition at the end of the expiratory phase. This combination of phasic excitatory drive and disinhibition supports their active phase during inspiration. Interestingly, we observed some concurrent phasic inhibition with the excitation during the I phase (see Figure 5 for higher temporal resolution conductance profiles from current-clamp recordings, and Figure 6 below incorporating conductances extracted from both current- and voltage-clamp data). Crucially, we did not observe discernible patterns of multi-phasic excitatory drive during the expiratory phase, and only steady levels of tonic inhibition predominated in this inter-burst interval. In the majority of recorded cells, the tonic inhibitory conductance during the expiratory phase exceeded any tonic excitatory conductance, functionally shunting the membrane potential.

**Figure 5:**
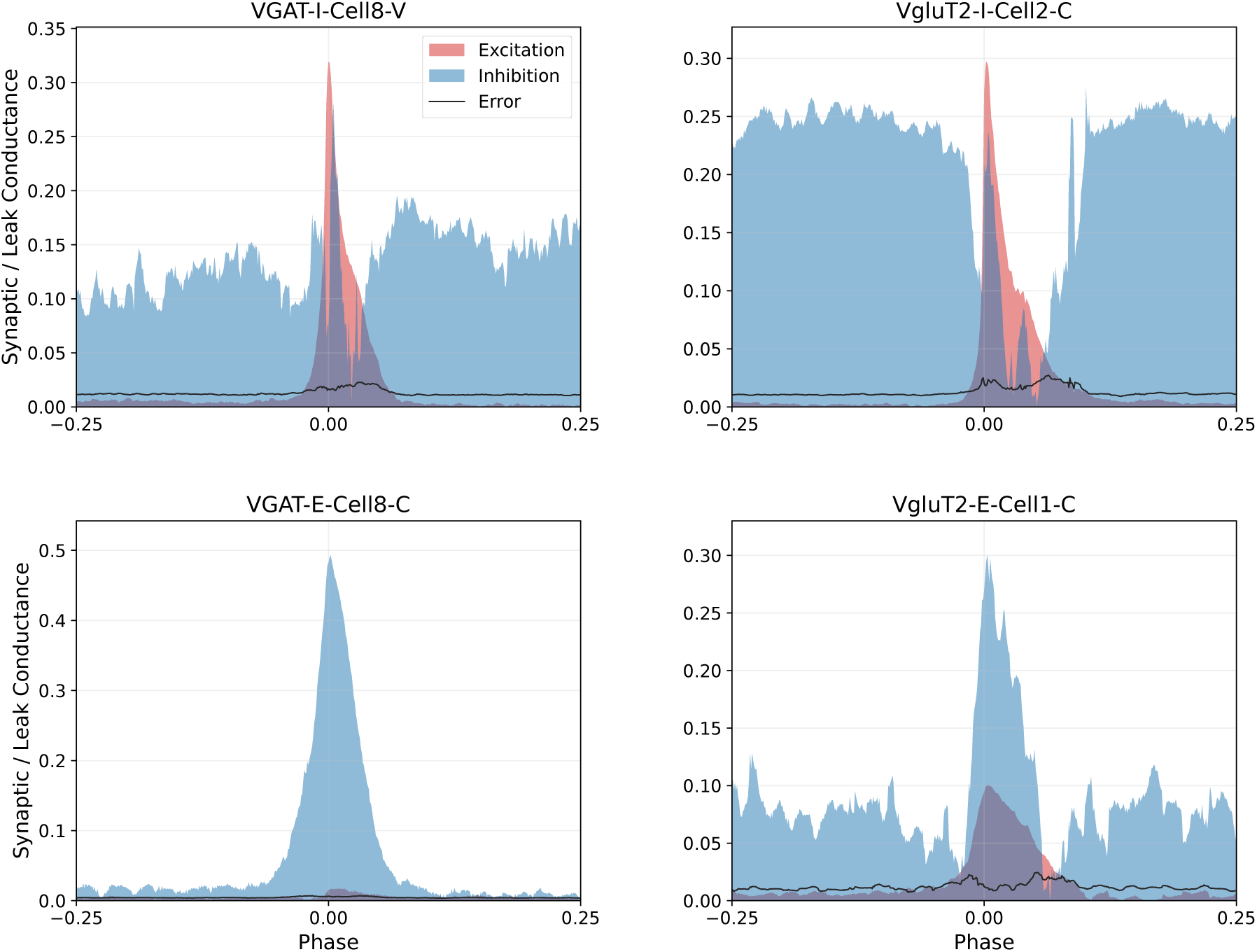
**Higher resolution phase-aligned reconstructed conductances for representative individual cells from each population group (VGAT-I, VgluT2-I, VGAT-E, VgluT2-E)**. Traces show excitatory (red) and inhibitory (blue) synaptic conductances normalized by the resting leak conductance, plotted over the normalized phase interval [−0.25, 0.25] surrounding the onset of the dominant burst (Phase 0). Solid black outlines indicate the standard error of the mean (SEM).

**Figure 6:**
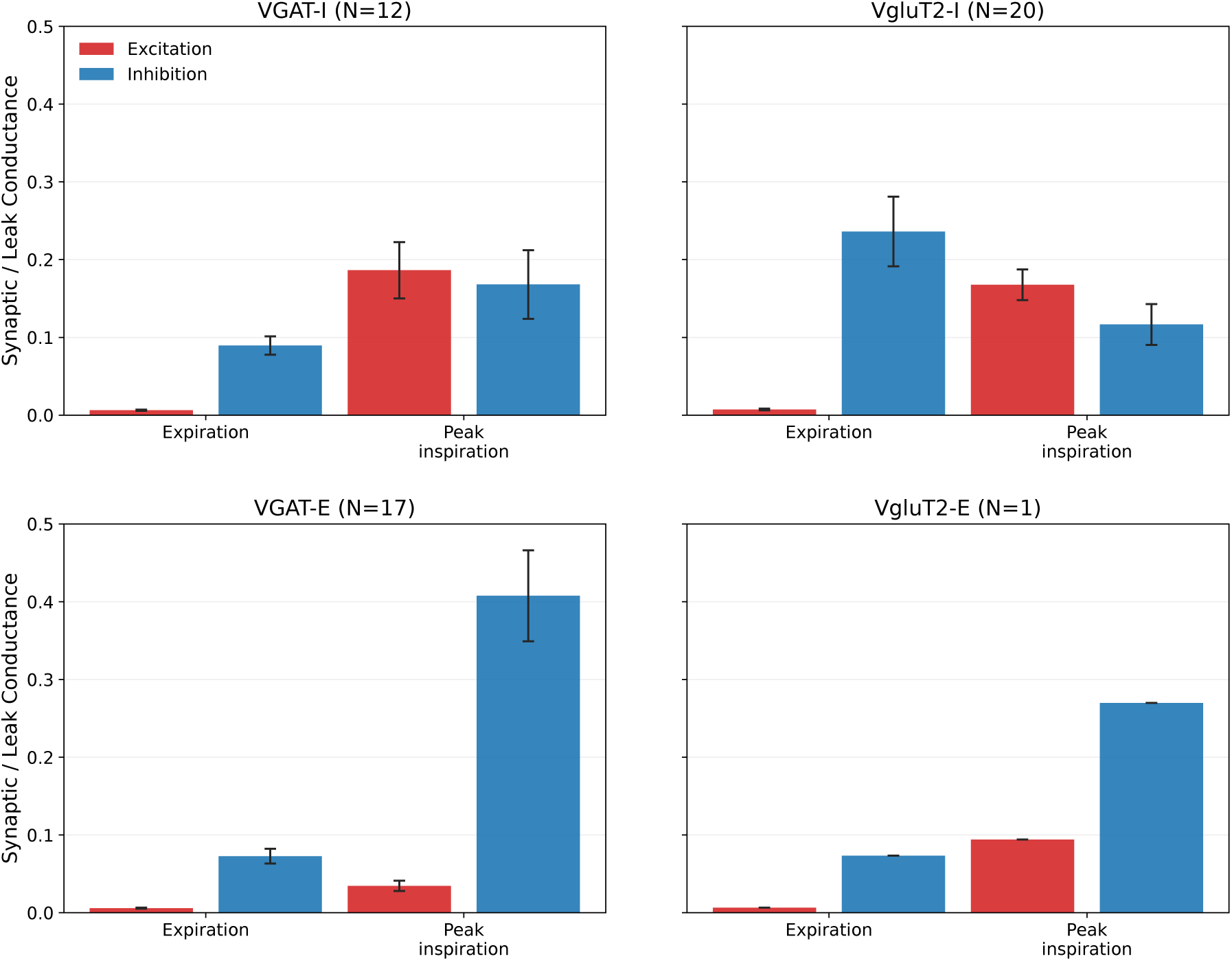
**Synaptic conductances during expiratory and inspiratory phases across respiratory neuron populations**. Bar graphs display the population mean (± SEM) normalized excitatory (*G*_exc_*/g*_leak_, red) and inhibitory (*G*_inh_*/g*_leak_, blue) conductances for VGAT-I (top left), VgluT2-I (top right), VGAT-E (bottom left), and VgluT2-E (bottom right) neurons. Conductances were summarized for the stationary inter-burst interval (mean expiratory value) and for the onset-aligned estimate of peak inspiratory conductance. The bars summarize the robust inlier subset after Tukey filtering of extreme values. The sample size (N) indicated for each group represents the number of analyzed recordings (each containing >25 respiratory cycles) rather than the number of unique cells, as some neurons contributed multiple recordings (both current- and voltage-clamp).

This dominance of tonic expiratory phase inhibition of inspiratory neurons provides a mechanistic explanation for the observed voltage trajectories (Figure 3): we consistently observed a smooth profile of developing excitation, with no evidence of regularly occurring burstlets or low-amplitude oscillations. This indicates that in the preBötC circuits isolated in the slice preparation, voltage trajectories and the suppression of ectopic burst spiking appear to be largely controlled by this potent synaptic inhibition.

Conversely, expiratory neurons exhibited a distinct pattern of synaptic inputs. They received strong phasic inhibition during the inspiratory phase, which effectively shunted their activity. We encountered only a single VgluT2-expressing neuron with this expiratory firing phenotype in our sample. This rare VgluT2-E neuron received no significant excitatory or inhibitory inputs during the expiratory phase, suggesting that the steady, low-frequency tonic firing of these neurons is potentially intrinsic or driven by tonic baseline cellular properties rather than by specific synaptic drive.

Critically, these synaptic input patterns appear to define the firing phenotype (inspiratory vs. expiratory) independent of the neuron’s own neurotransmitter identity. Both VgluT2+ and VGAT+ neurons within the same firing class shared identical synaptic input profiles.

### Neuron-Type and Phase-Specific Normalized Synaptic Conductances

To decipher the effective synaptic connectivity between the key active respiratory populations, we analyzed the magnitude of excitatory and inhibitory conductances during estimated peak inspiration and during the stationary expiratory/interburst interval (Figure 2). Because individual neurons vary in physical size, and since both membrane capacitance and resting leak conductance scale proportionally with cell surface area (Koizumi et al., 2008), we normalized all synaptic conductances to each neuron’s respective leak conductance. This size-independent normalization controls for morphological variability, providing a standardized metric of the effective synaptic drive relative to the cell’s intrinsic passive properties. Based on the phase-specificity of activity, we attribute inhibition during inspiration to VGAT-I neurons and inhibition during expiration to VGAT-E neurons. Excitation is attributed to VgluT2-I neurons and is concentrated at estimated peak inspiration, while excitatory inputs during expiration were found to be negligible.

The relative magnitudes of the synaptic conductances for each population are detailed in **Table 2** and visualized as population means in Figure 6.

**Table 2:** Summary of reconstructed synaptic conductances across neuronal populations. Population mean ± SEM of leak-normalized excitatory (*G*_exc_*/g*_leak_) and inhibitory (*G*_inh_*/g*_leak_) conductances for the onset-aligned estimate of peak inspiration and for the stationary inter-burst (expiratory) interval. *N* denotes the number of recordings retained after Tukey IQR outlier filtering applied per group and per metric to the cycle-count-screened set in Table 1, and may therefore differ from the corresponding row of Table 1; current- and voltage-clamp recordings are pooled. SEM is reported as N/A for the VgluT2-E group, which is based on a single recording.

| Group (N) | Phase | $G_{\text{exc}}/g_{\text{leak}}$ | $G_{\text{inh}}/g_{\text{leak}}$ |
| --- | --- | --- | --- |
| VGAT-I (12) | Expiration | $0.007 \pm 0.001$ | $0.087 \pm 0.012$ |
| | Peak inspiration | $0.186 \pm 0.036$ | $0.167 \pm 0.044$ |
| VgluT2-I (18) | Expiration | $0.007 \pm 0.001$ | $0.236 \pm 0.045$ |
| | Peak inspiration | $0.168 \pm 0.020$ | $0.117 \pm 0.026$ |
| VGAT-E (17) | Expiration | $0.006 \pm 0.001$ | $0.073 \pm 0.009$ |
| | Peak inspiration | $0.035 \pm 0.007$ | $0.408 \pm 0.058$ |
| VgluT2-E (1) | Expiration | $0.007 \pm \text{N/A}$ | $0.073 \pm \text{N/A}$ |
| | Peak inspiration | $0.094 \pm \text{N/A}$ | $0.270 \pm \text{N/A}$ |

**VGAT-I Population.** In the top-left panel, these inhibitory inspiratory neurons received multiple phase-locked synaptic inputs. Their largest measured conductance was excitatory input during the inspiratory phase, accompanied by substantial concurrent inhibition during inspiration. Inhibitory conductance during expiration was also present but of a smaller magnitude.

**VgluT2-I Population.** In the top-right panel, the largest conductance measured in the VgluT2-I population was inhibition during the expiratory phase. During inspiration, these neurons exhibited considerable excitatory conductance, along with a smaller degree of concurrent inhibition.

**VGAT-E Population.** In the bottom-left panel, the synaptic profile of inhibitory expiratory neurons was dominated by inhibition during the inspiratory phase, which was the largest conductance observed for this group. During their active expiratory phase, they exhibited a much smaller degree of inhibition. Excitatory conductance during the inspiratory phase was present but minimal.

**VgluT2-E Population.** Although only a single VgluT2-E neuron was recorded, its pattern of synaptic conductances was largely similar to that of the VGAT-E population. In the bottom-right panel, its largest measured conductance was inhibition during the inspiratory phase, accompanied by a smaller excitatory conductance during inspiration. During its active expiratory phase, inhibitory conductance was minimal and excitatory conductance was negligible.

In summary, the synaptic conductance profiles in Figure 6 and Table 2 reveal prominent, phase-locked inhibitory inputs across both phases, with the major excitatory inputs restricted primarily to the inspiratory phase.

### Inferred Circuit Functional Connectivity

Based on the measured synaptic conductance profiles, we can infer the specific functional connectivity between the recorded populations (Figure 7).

**Figure 7:**
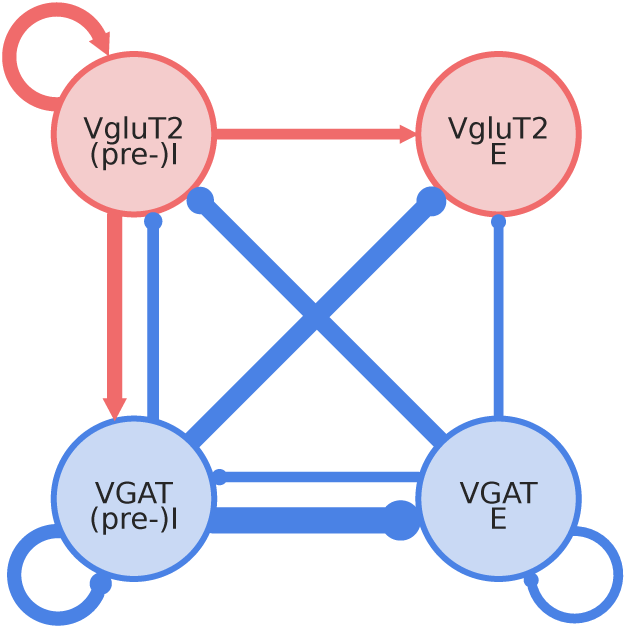
**Conductance-weighted circuit connectivity diagram**. Inferred preBötC population circuit with connection line widths scaled to the same mean normalized conductances summarized in Table 2. Red arrows indicate excitatory drive from the inspiratory VgluT2 population, blue lines with terminal circles indicate inhibitory drive from the inspiratory and expiratory VGAT populations, and only connections exceeding the display threshold are shown. Neurons with pre-I spiking (pre-) are included as constituents of the VgluT2-I and VGAT-I populations. Connections below 0.05 are clipped from the diagram.

Here and below, terms such as recurrent excitation, recurrent inhibition, self-excitation, and self-inhibition are used at the level of functional populations. They denote inferred interactions among neurons within the same genetically and phase-defined population, such as VgluT2-I to VgluT2-I or VGAT-I to VGAT-I, rather than autaptic feedback onto the same recorded neuron or direct proof of monosynaptic self-connections.

First, all populations receive inhibition during the expiratory phase, which is most consistent with input from the concurrently active inhibitory population: the VGAT-E neurons. Thus, we infer that VGAT-E neurons connect throughout, providing robust functional inhibition to the VgluT2-I population, along with weaker inhibitory inputs to the VGAT-I and VgluT2-E populations, as well as recurrent inhibition onto themselves (Figure 7).

Second, all populations also receive prominent inhibition during the inspiratory phase. This inhibition must arise from the inhibitory population active during inspiration: the VGAT-I neurons. This indicates a functional inhibitory network originating from VGAT-I, connected with high conductance magnitude to both expiratory populations (VgluT2-E and VGAT-E), while also providing substantial input to the VgluT2-I population and forming a robust recurrent inhibitory loop within the VGAT-I population itself (Figure 7).

Third, the phasic excitation received during the inspiratory phase must be driven by the excitatory population active at that time: the VgluT2-I neurons. This pattern is consistent with prominent recurrent excitation within the VgluT2-I population, as well as robust feed-forward excitation to the VGAT-I and VgluT2-E populations (Figure 7). Excitatory input to the VGAT-E population was below the threshold for representation in the circuit schematic.

Finally, the lack of significant excitatory synaptic drive received by any population during the expiratory phase specifically excludes prominent recurrent excitation within the VgluT2-E population, as well as feed-forward excitation from VgluT2-E to any other recorded population in this slice preparation. Therefore, we do not infer functional excitatory outputs originating from the VgluT2-E population during baseline in vitro rhythmic activity. Although we identified only a single VgluT2-E neuron in our sample of recorded neurons, we postulate that this type of neuron represents a functional class and therefore needs to be recorded more extensively to definitively determine their functional role.

These assignments represent functional attributions based on phase relationships and do not exclude contributions from additional sources or polysynaptic pathways.

## Discussion

### Circuit Organization Intrinsic to the preBötC and the Rhythmogenic Kernel

While the preBötC has long been recognized as the primary site of inspiratory rhythm generation, the organization of its core excitatory–inhibitory circuitry has not been established. Many experimental and theoretical analyses have focused on rhythmogenic mechanisms operating under in vitro conditions in rhythmically active slices, providing insights into the biophysical and circuit processes involved. As discussed in many articles since the slice preparation was developed (Smith et al., 1991), these insights are relevant for understanding rhythm and pattern generation under various conditions in vivo (e.g., Smith et al., 2000; Feldman and Del Negro, 2006; Smith et al., 2007; Richter and Smith, 2014; Phillips and Rubin, 2019). Previous studies have established patterns of inspiratory and expiratory neuron activity in the preBötC in vitro (e.g., Johnson et al., 1994; Shao and Feldman, 1997; Carroll et al., 2013), which also exist in more intact en bloc in vitro preparations (Smith et al., 1990). Within these electrophysiological phenotypes, the subpopulations of rhythmically active excitatory and inhibitory neurons (e.g., Kuwana et al., 2006; Winter et al., 2009; Morgado-Valle et al., 2010; Koizumi et al., 2013; Ausborn et al., 2018) and their synaptic interactions have not been thoroughly characterized. Therefore, any analysis of the functional circuit organization intrinsic to the preBötC must account for the synaptic interactions among these distinct interneuron types and explain how such interactions produce neuronal firing patterns and contribute to rhythm generation. The structural-functional properties of preBötC excitatory and inhibitory populations have been previously analyzed in seminal studies (Koizumi et al., 2013). However, how these populations functionally interact to generate neuronal firing patterns and a robust rhythm remains unresolved. Our inference technique for synaptic conductances in rhythmic circuits (Molkov et al., 2025), which we have applied in the present studies, provides a novel quantitative view of this functional organization of preBötC circuits active in slices. Below, we discuss features of this organization within the framework of the rhythmogenic “kernel” proposed in earlier experimental and modeling studies. We also consider how our inferred in vitro circuit functional connectome relates to the connectome we have found for preBötC circuits in more intact in situ states.

#### Functional Features of the Circuit Organization

The excitatory and inhibitory synaptic interactions as described in Results are summarized by the conductance-weighted circuit schematic in Figure 7, which represents the functional connectome of the active neuronal populations in vitro. The synaptic conductances extracted from our analyses and represented in Figure 7 reflect the total set of synaptic inputs arising from the active neuronal populations indicated. This schematic highlights the central role of synaptic interactions in shaping neuronal activity. The self-excitatory VgluT2-I population forms the rhythm-generating kernel, where strong recurrent excitation synchronizes and drives rhythmic inspiratory bursts. There is also concurrent dynamic inhibition and excitation in the VgluT2-I and VGAT-I populations. This kernel is regulated by its interconnections with a robust, reciprocal inhibitory scaffold (VGAT-I ↔ VGAT-E), a main component of the inhibitory connectome that enforces phase-locking and defines the inter-burst interval. Figure 7 provides the first detailed depiction of the functional inhibitory connectome in the preBötC in vitro, revealing greater complexity than previously recognized.

The dynamic interactions implied by Figure 7 are consistent with the view that respiratory rhythm generation is dynamically assembled from interacting microcircuits whose synchronization depends on the balance of recurrent excitation, intrinsic excitability, and concurrent inhibition (Ramirez and Baertsch, 2018). Our conductance reconstructions provide a direct synaptic-level description of this balance in the isolated rhythmic slice state.

Another feature of this connectome is that the VgluT2-I and VGAT-I populations contain neurons with pre-I spiking patterns, an established feature of preBötC neuronal activity (e.g., Johnson et al., 1994; Butera et al., 1999b; Del Negro et al., 2001). However, this activity has not previously been identified for inhibitory neurons in vitro. The presence of pre-I activity in both VgluT2+ and VGAT+ populations indicates that the initiation of the inspiratory phase involves the early, simultaneous recruitment of both excitatory and inhibitory neuron classes. In this network, pre-I excitatory neurons likely provide early synaptic drive to gradually recruit the excitatory rhythmogenic kernel and the corresponding inhibitory interneurons. We suggest this pre-I activity supports a specific mechanistic sequence for the expiratory-to-inspiratory phase transition in the slice circuits. Early-activating pre-I excitatory neurons drive gradual network recruitment, including pre-I inhibitory neurons. These inhibitory pre-I neurons then provide early synaptic inhibition needed to suppress the expiratory population (VGAT-E), which disinhibits inspiratory populations and facilitates the phase transition. The functional consequence of early inhibitory recruitment is shown by voltage and synaptic current profiles of expiratory cells during the late expiratory phase (Supplemental Figure 4). Both VGAT-E and VgluT2-E neurons stop firing before the global inspiratory burst. Matched voltage-clamp recordings show that this pre-inspiratory silent window coincides with the early onset of outward (inhibitory) synaptic currents. This developing pre-I inhibition effectively shunts and ends expiratory spiking before the main inspiratory burst emerges. The coordinated pre-I inhibitory and excitatory neuronal activity appears to be a fundamental feature of dynamic circuit operation, with neuronal synaptic interactions contributing to phase switching alongside other inhibitory interactions depicted in Figure 7.

As indicated above, the inhibitory connectome also features the feed-back interaction of VGAT-I to the VgluT2-I excitatory kernel. Thus, during the generation of the inspiratory population burst, there is concurrent synaptic excitation and a smaller phasic inhibition in the VgluT2-I population (Figures 5 and 6). This phasic inhibition likely dynamically shapes the temporal evolution of the inspiratory burst and possibly regulates its amplitude. Pharmacological block of synaptic inhibition in the isolated preBötC has previously been shown to augment inspiratory population burst amplitude (e.g., (Johnson et al., 2001)).

Overall, our circuit analyses based on synaptic conductance profiling indicate that, beyond rhythm generation, the circuits intrinsic to the preBötC are designed to generate coordinated patterns of inspiratory and expiratory activity through interactions between an excitatory rhythmogenic kernel and a functional inhibitory connectome. The latter has complex connectivity within the network.

#### The VgluT2-(pre-)I Rhythmogenic Kernel

Classic experimental and modeling studies have proposed—and there is general agreement—that the preBötC excitatory circuits have intrinsic autorhythmic properties. When isolated in slices in vitro under appropriate excitability conditions, these circuits continue to intrinsically generate rhythmic activity that drives behaviorally relevant inspiratory hypoglossal motoneuronal output. Experiments also showed that rhythmic inspiratory population activity can be maintained within a physically isolated preBötC region (‘preBötC island’) and within a hemi-slice, even after blocking fast synaptic inhibition (GABAergic and glycinergic). This indicates that a minimal, local excitatory circuit, relying mainly on synaptic coupling and recurrent excitation to recruit and synchronize neuron activity, is sufficient to generate inspiratory population bursts and rhythm (Del Negro et al., 2001; Johnson et al., 2001). Thus, the essential rhythmogenic kernel consists of a self-excitatory population of synaptically coupled glutamatergic neurons, as depicted in Figure 7. This concept is supported by many experimental and modeling studies (Del Negro et al., 2001; Johnson et al., 2001; Smith et al., 2007; Rubin et al., 2009; Molkov et al., 2017; Phillips and Rubin, 2019; Phillips et al., 2022; Ramirez and Baertsch, 2018), though there is no consensus about the underlying rhythmogenic mechanism(s) (Butera et al., 1999a,b; Del Negro et al., 2018; Phillips and Rubin, 2019; Phillips et al., 2022). One key property of the rhythmogenic mechanism is the voltage-dependence of bursting and its frequency control over a wide dynamic range (Butera et al., 1999a,b; Del Negro et al., 2001; Koizumi et al., 2013; Phillips et al., 2022). The failure of inhibitory synaptic blockade to disrupt in vitro rhythm generation means that the inhibitory connectome in Figure 7 is not required for inspiratory rhythm generation in vitro. However, the excitatory kernel’s dynamical operation is apparently modulated by the level and structure of inhibition. Instabilities of the rhythm—cycle-to-cycle variability of period and amplitude—and changes of excitability, including voltage-dependence of frequency control of the rhythm-generating kernel, occur after blocking synaptic inhibition in vitro (Johnson et al., 2001). These changes align with ongoing synaptic inhibition from VGAT-E neurons and inhibitory connections from VGAT-I, which regulate the excitability of the VgluT2 kernel.

These earlier studies noted above established the concept of a rhythmogenic kernel consisting of a population of excitatory neurons with intrinsic rhythmogenic properties, synaptically coupled by local excitatory amino acid (non-NMDA)-mediated synaptic interactions (Koshiya and Smith, 1999). Morphological studies in the rhythmic slice demonstrated that a majority of preBötC excitatory inspiratory neurons have contralaterally projecting axons, enabling left-right inspiratory population burst synchronization in the bilateral preBötC, and that a subset of these commissural excitatory neurons has intrinsic oscillatory bursting properties, which was proposed to be an important property of the excitatory kernel neurons (Koizumi et al., 2013). These studies also indicated that preBötC inhibitory interneurons generally lack intrinsic rhythmic bursting properties and contralateral axonal projections, and therefore mainly interact locally within the preBötC. Our analyses do not identify the sources of synaptic inputs, but the morphological reconstruction results imply that the excitatory conductances we infer can arise from both local and bilateral excitatory connections, whereas the inhibitory conductances arise from local synaptic interactions. Our conductance-resolved mapping of synaptic interactions in the preBötC in vitro builds directly on the foundational structural-functional characterization of excitatory and inhibitory respiratory interneurons by Koizumi et al. (2013) by delineating the functional inhibitory synaptic interactions with the excitatory kernel.

Recent experimental and modeling studies (Phillips and Rubin, 2019; Phillips et al., 2022) also support the notion that a subpopulation of preBötC excitatory neurons serves as the kernel for rhythm generation. A VgluT2-I subpopulation in the preBötC amplifies synaptic drive from the kernel population to generate the amplitude of the inspiratory population burst. This implies that the excitatory neurons active in the slice have a more complicated functional organization than depicted in earlier studies and Figure 7. We suggest that both of these functionally distinct subpopulations of VgluT2-I neurons are subject to the set of inhibitory interactions depicted in Figure 7. This remains to be established, since we did not experimentally differentiate these functional types of excitatory neurons by protocols to identify cells with intrinsic, voltage-dependent rhythmic bursting properties (e.g., Koizumi et al., 2013), nor did we specifically determine their synaptic conductance profiles.

Despite this complexity, optogenetic photostimulation of the whole VgluT2 population in vitro produces the same voltage-dependent regulation of inspiratory burst frequency (Phillips et al., 2022) predicted for the rhythmogenic subpopulation. This observation suggests that the excitatory synaptic interactions among sub-populations define the behavior of the whole population. Thus, the self-excitatory VgluT2-(pre-)I population depicted in Figure 7 can serve as a convenient representation of this behavior.

Together with the foundational (Johnson et al., 1994, 2001; Koizumi et al., 2013) and the more recent studies, the present circuit analysis supports the interpretation that the preBötC in the slice preserves a functional excitatory rhythmogenic kernel best understood as a bilaterally distributed excitatory microcircuit whose dynamics are stabilized by tonic expiratory phase inhibition that regulates the interval between inspiratory bursts, and phasic inhibitory interactions involved in coordinating phase transitions.

### Preserved Core Circuitry and Reduced Functional Inhibitory Circuit Interactions in the Isolated preBötC

Our main finding is that the functionally active classes of preBötC expiratory and inspiratory neuron populations, as well as inhibitory interactions, are reduced in the in vitro slice compared to how the preBötC operates in a more intact brainstem. Direct comparison with our previous conductance-based circuit reconstructions for the in situ respiratory network (Molkov et al., 2025, Figure 8 below) highlights this simplification: the in situ preBötC shows multiphasic excitatory and inhibitory interactions, involving post-inspiratory and augmenting expiratory neuron activity. In contrast, the slice exhibits a simplified pattern—dominated by tonic inhibition from a population of expiratory neurons (VGAT-E) and phasic (pre-)inspiratory inhibition from VGAT-(pre-)I neurons. This indicates that the “reduced” slice is simplified functionally, not just anatomically, in its synaptic interactions and pattern generation mechanisms. How does the preBötC connectome in vitro relate to the functional connectome operating in more intact conditions?

**Figure 8:**
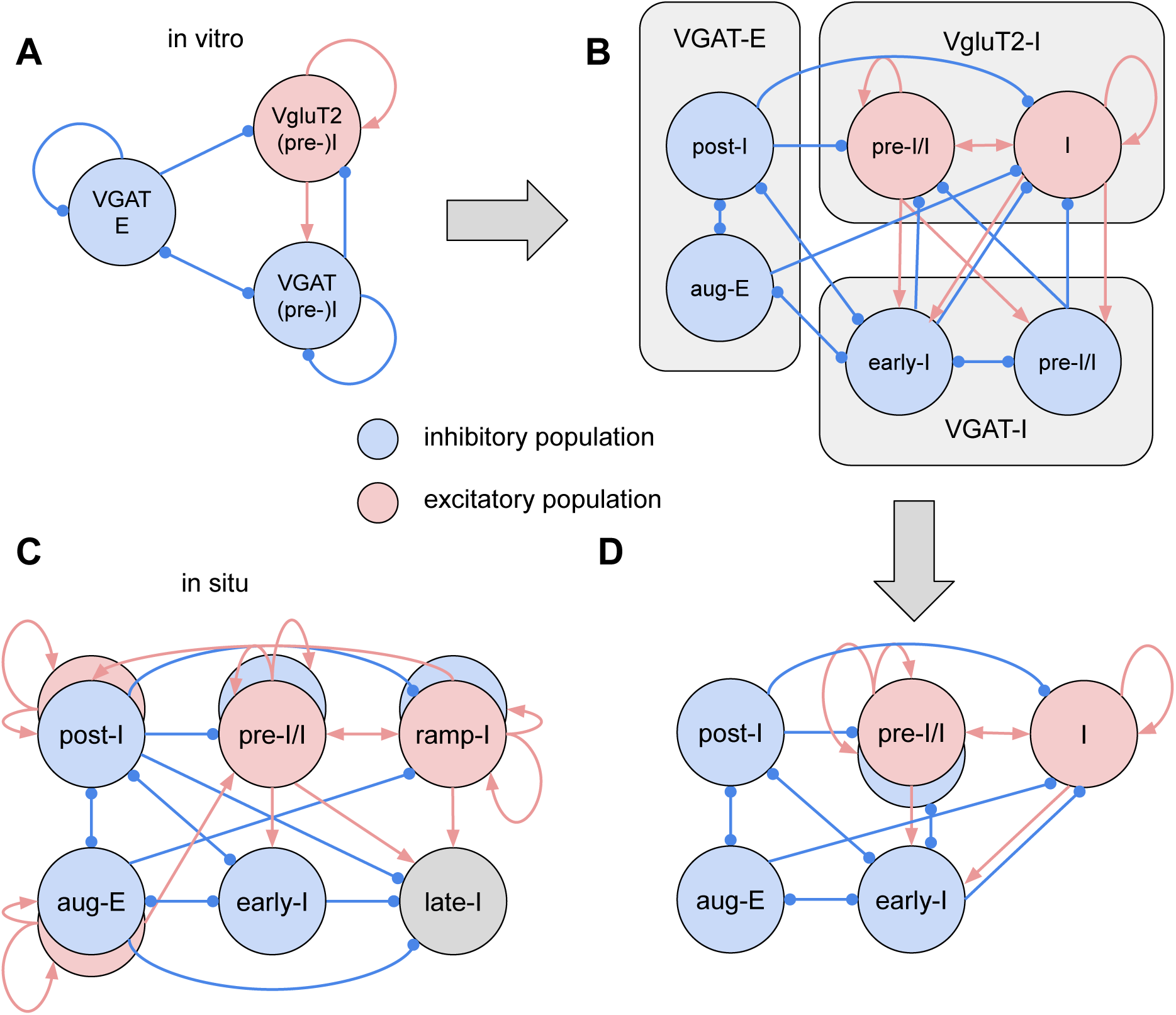
**Preserved core circuitry and reduced functional inhibitory architecture in the isolated preBötC in vitro**. (A) Compact in vitro circuit derived from our conductance analyses for the rhythmic slice. (B) Interpretive resolution of modest intra-population conductances, attributed to self-inhibition in A, into weak cross-inhibition between temporally offset subsets, with tentative mapping onto classic in situ neuronal classes for the active preBötC neuron types. (C) Full in situ connectome (Molkov et al., 2025) showing the expanded inhibitory connectome. (D) Simplified core circuit interactions that may be preserved in the slice, consisting of a reciprocal inhibitory circuit motif with pre-I/I coordination. Red arrows indicate excitatory connections; blue arrows indicate inhibitory connections. See also Supplemental Figures 3–4 for pre-I dynamics relevant to this interpretation.

Classical models (e.g., Smith et al., 2007; Rubin et al., 2009; Lindsey et al., 2012; Richter and Smith, 2014) describe how the preBötC operates within the ventral respiratory column and outline a hierarchical organization. In these models, the preBötC generates a basic inspiratory rhythm via an excitatory interneuron kernel, including pre-I spiking neurons. PreBötC inhibitory inspiratory neurons suppress expiratory activity and coordinate inspiratory-expiratory transitions. To produce the full three-phase respiratory pattern—(pre-)inspiratory, post-inspiratory (post-I), and late (aug-E) expiratory activity—which is observed in the intact brainstem, additional inhibitory populations such as post-I and aug-E neurons from the rostral Bötzinger Complex (BötC) are required. Large-scale computational models propose that this three-phase pattern depends on a “mutually inhibitory ring” structure linking the preBötC and BötC inhibitory circuits (Smith et al., 2007; Rubin et al., 2009; Molkov et al., 2017). Phase transitions are thus controlled by reciprocal inhibition between preBötC inspiratory (early-I) and BötC expiratory (post-I and aug-E) inhibitory neurons. For more detailed connectome comparisons, consult prior models and schematics from our conductance analyses (Figure 8B; Supplemental Figure 1 in Molkov et al. (2025)).

Recent experimental and modeling studies have revealed a more complex inhibitory connectome within the preBötC, including post-I inhibitory neurons and their interactions with excitatory and inhibitory inspiratory neurons, which were absent from earlier models. Consistent with prior in vivo studies (e.g., Schwarzacher et al., 1995), our in situ recordings (Molkov et al., 2025) identified substantial numbers of post-I and aug-E neurons, including glycinergic inhibitory subtypes within the preBötC. We also observed diverse inspiratory activity patterns and synaptic conductances, and neurons identified in situ display multiphasic synaptic interactions aligned with three-phase pattern generation. Thus, preBötC neurons—including inhibitory types—generate various inspiratory and expiratory activity patterns within a three-phase organization. These neurons with distinct firing phenotypes in situ are not fully engaged in the isolated, rhythmically active medullary slices, where neuronal activity patterns are transformed.

Our conductance-based in vitro and in situ preBötC circuit reconstructions (Figures 7 and 8; Molkov et al., 2025) indicate that the isolated preBötC contains core circuits, including the inhibitory connectome, capable of generating both inspiratory and expiratory patterns. Notably, certain connectivity motifs observed in vitro resemble those found in the intact network. The main motifs retained include excitatory and inhibitory populations for each firing phenotype [(pre-)I and E], as well as reciprocal interactions between I and E inhibitory neurons.

The VgluT2-E population shown in Figure 7 is not carried forward as a distinct node in Figure 8 because Figure 7 summarizes conductance inputs to all recorded firing classes, whereas Figure 8 is an interpretive comparison focused on the preserved rhythmogenic excitatory kernel and inhibitory circuit motifs. Since VgluT2-E neurons were sparsely sampled and were not inferred to provide functional excitatory output during baseline slice rhythmic activity, we do not assign this population a separate role in the reduced core schematic.

In Figure 8 we attempt to specifically relate the in vitro (Figure 8A) and in situ (Figure 8C) functional connectomes. We consider that the modest inhibitory conductances attributed (Figure 7) to self-inhibition (VGAT-I ≈ 0.167 during inspiration; VGAT-E ≈ 0.073 during expiration) may instead reflect weak cross-inhibition within temporally offset neuronal subsets, not true self-inhibition (Figure 8B). In this view, the VGAT-I population divides into pre-I/I and early-I inhibitory subsets (using the phenotype nomenclature for in situ preBötC activity patterns) with moderate reciprocal interactions, while the VGAT-E population divides into post-I and aug-E inhibitory subsets. These coordinated subsets become active once the preBötC operates in the ventral respiratory column in situ or in vivo. Although inferential, this mapping suggests clear population-level parallels between the slice-derived circuit and the in situ functional neuronal classes and their connectivity.

This interpretation is consistent with the neuronal firing patterns and synaptic conductance profiles in the preBötC operating within the intact ventral respiratory column. Earlier models (e.g., (Smith et al., 2007)) placed post-I and aug-E populations primarily in the more rostral BötC circuits for expiratory pattern generation. However, our analysis of synaptic conductance profiles shows that the active preBötC post-I and aug-E neurons in situ have profiles similar to those of their BötC counterparts. We do not currently know if the preBötC circuits can intrinsically generate these expiratory patterns under the appropriate excitability conditions without external inputs. Alternatively, inputs—including those from the BötC—seem to be required to interact with specific subsets of preBötC expiratory neurons. When the preBötC and BötC operate together in situ without pontine inputs, their interacting circuits generate only a two-phase inspiratory-expiratory rhythmic pattern (Smith et al., 2007). Thus, while the BötC circuits are clearly connected to preBötC circuits—including inhibitory circuit interactions (Ausborn et al., 2018) that are generally required for three-phase pattern generation (Richter and Smith, 2014; Marchenko et al., 2016)— the BötC circuits are not the exclusive source of these patterns. However, they are important for elaborating these patterns as observed in situ (Figure 8C). Notably, pharmacological disruption of synaptic inhibition in the preBötC in situ disrupts the three-phase respiratory pattern and causes pronounced disturbances in the rhythm (Marchenko et al., 2016). Likewise, disrupting inhibition in the BötC disrupts this pattern and perturbs the rhythm. The interactions of preBötC and BötC circuits apparently reinforce each other in generating the three-phase rhythmic pattern. As noted earlier, disrupting synaptic inhibition in the preBötC in vitro does not disrupt rhythm generation. From this perspective, the slice preserves a compact core inhibitory circuit motif (Figure 8A), but lacks the extended circuitry with a more complex inhibitory and excitatory connectome that is normally required for three-phase pattern generation.

This framework, outlined in Figure 8, is further supported by analogous pre-inspiratory dynamics (Supplemental Figures 3–4). Specifically, the inhibitory pre-I/I population, present in both in situ and in vitro (Supplemental Figure 3), functions as an underappreciated but important component within the preBötC circuits by providing inhibitory input to inspiratory neurons. Earlier computational models (Smith et al., 2007; Ausborn et al., 2018; Butera et al., 1999b) typically emphasized inspiratory phase-switching mechanisms involving pre-I excitatory drive (serving as an inspiratory on-switch) and post-I inhibition (acting as an off-switch). The inhibitory pre-I/I population may participate in terminating the expiratory phase and in organizing the timing and balance of activity among inspiratory inhibitory and excitatory populations within the preBötC circuit.

In summary, the slice isolates a compact and functionally reduced version of the in situ network. In this sense, the slice does not simply remove anatomical components; it reveals a different functional configuration of the respiratory network, converting the richer multiphasic organization of the intact system into a reduced two-phase pattern. Specifically, we hypothesize this reduction—from synaptic interactions producing a three-phase rhythmic pattern to interactions only generating a two-phase alternating rhythmic pattern—results from the lack of excitatory drive from pontine and perhaps retrotrapezoid nuclei and the absence of BötC circuit interactions and postinspiratory complex (PiCo) neuron inputs (Anderson et al., 2016) in the isolated slice preparation. In the intact network, higher baseline excitability and neuronal spike-frequency adaptation enable embedded subpopulations to activate, thereby driving sequential multiphasic transitions. In the slice, without these external drives, reduced excitability limits transitions. Consequently, the preBötC’s intrinsic excitatory neurons generate a rhythm that the preserved reciprocal inhibitory components convert into a binary alternating pattern. Thus, external excitatory and BötC inputs expand the existing preBötC circuit into richer multiphasic inhibitory sequences and expiratory activity. The isolated preBötC retains core reciprocal inhibitory circuit organization and pre-I/I excitatory–inhibitory coordination, but lacks the complex inhibitory and excitatory interconnections present in the intact network.

### Limitations and Developmental Considerations

We have discussed potential limitations of our conductance analysis methods and assumptions in our previous publication (Molkov et al., 2025). We also presented experimental validation based on current- and voltage-clamp recordings with sharp electrodes in situ. For our in vitro whole-cell patch-clamp recordings, we repeated experimental validation and addressed the main issues (see Supplemental Materials).

We targeted excitatory and inhibitory preBötC neurons to identify the main active functional cell classes. As noted above, excitatory preBötC neurons have distinct functional subtypes, including inspiratory neurons that drive rhythm generation and neurons that control inspiratory burst amplitude (Phillips and Rubin, 2019; Phillips et al., 2022). The excitatory inspiratory populations vary in intrinsic properties, burst firing patterns, and axonal projections (Koizumi et al., 2013). Inhibitory neurons may also have diverse subtypes based on neurotransmitter phenotype and axonal projections (Koizumi et al., 2013) with distinct roles. We may have recorded only from some of these subsets, thereby missing important subpopulations and their synaptic interactions. The excitatory expiratory neurons represented in Figure 7 are an example of an undersampled neuronal type in our recordings, and, as mentioned above, require more functional analysis. Thus the functional connectome represented in Figure 7 may be overly simplified; more elaborate interpretations of circuit functional architecture are possible (Figure 8B). Nevertheless, the consistent synaptic conductance profiles across our recorded cell classes suggests that at least the basic structure of synaptic interactions— phasic drives and tonic expiratory inhibition—is a robust feature of the in vitro active network, not a result of sampling bias.

We have attempted to understand transformations in neuronal activity patterns and synaptic interactions in vitro versus intact in situ conditions. However, this comparison has clear limitations, mainly related to developmental and excitatory states. Our slice data came from neonatal (P3-P8) mice, while in situ data were from older juveniles (3-4 weeks old rats). Elevated K^+^ (8-9 mM) is required to sustain rhythmic activity in slices, and as noted earlier, the lack of external inputs creates an excitation state different from that in the in situ arterially perfused preparation, which also has more normal K^+^ levels. As shown (Smith et al., 2007), removing brainstem structures rostral to the preBötC in situ and elevating K^+^ to more closely match slice excitability produce inspiratory patterns and rhythms similar to those of the neonatal slice, showing that developmental state is not an issue. These findings highlight the crucial role of external inputs from circuits rostral to the preBötC in three-phase rhythmic pattern generation and in changes in synaptic interactions in their absence.

## Conclusion

Our inferred functional preBötC circuits and synaptic interactions in vitro reveal a reduced functional state of the in vitro respiratory network. In this state, the complex multiphasic inhibition and excitation observed in more intact systems is replaced by a simpler excitatory and inhibitory circuit functional architecture. This synaptic organization features interactions between a self-exciting VgluT2 rhythmogenic kernel and an inhibitory connectome with more complex interactions than previously understood. The inhibitory connectome provides inspiratory phasic inhibition for phase switching and tonic expiratory inhibition for controlling the inspiratory interburst interval. We propose that this inhibitory control is an important circuit mechanism for regulating the VgluT2 population’s rhythmogenic mechanism in vitro. Our circuit analyses are important for interpreting mechanisms of rhythm and pattern generation in vitro. They provide a crucial framework for mapping state-dependent changes in circuit logic, bridging the gap between the reduced in vitro models and more complex intact systems.

## Ethics Statement

All animal procedures were approved by the Animal Care and Use Committee of the National Institute of Neurological Disorders and Stroke (Animal Study Proposal #1154-21).

## Acknowledgments

This work was supported by the Intramural Research Program of the NIH, NINDS.

## Code and Data Availability

All raw whole-cell patch-clamp recordings analyzed in this study, together with the C++ analyzer source code, Python processing and figure-generation scripts, per-recording configuration files, and processed conductance summaries, are publicly available at https://github.com/ymolkov/synaptic-inputs-slices under the Creative Commons Attribution 4.0 (CC BY 4.0) license. A companion site that renders the per-recording analysis artifacts as an interactive static dashboard is hosted at https://ymolkov.github.io/synaptic-inputs-slices/.

## Supplemental Materials

### Sensitivity Analysis of Reversal Potentials

The robustness of the synaptic conductance inference was evaluated by systematically varying the assumed reversal potentials for excitation (*E_e_*) and inhibition (*E_i_*) within biologically plausible ranges (Supplemental Figure 1). The resulting 3×3 matrix illustrates the reconstructed excitatory (*G_exc_*, red) and inhibitory (*G_inh_*, blue) conductances across combinations of *E_e_* at -20.0, -10.0, and 0.0 mV, and *E_i_* at -90.0, -80.0, and -70.0 mV. Each panel represents the mean conductance over two normalized respiratory cycles for a representative inspiratory neuron.

**Supplemental Figure 1.**
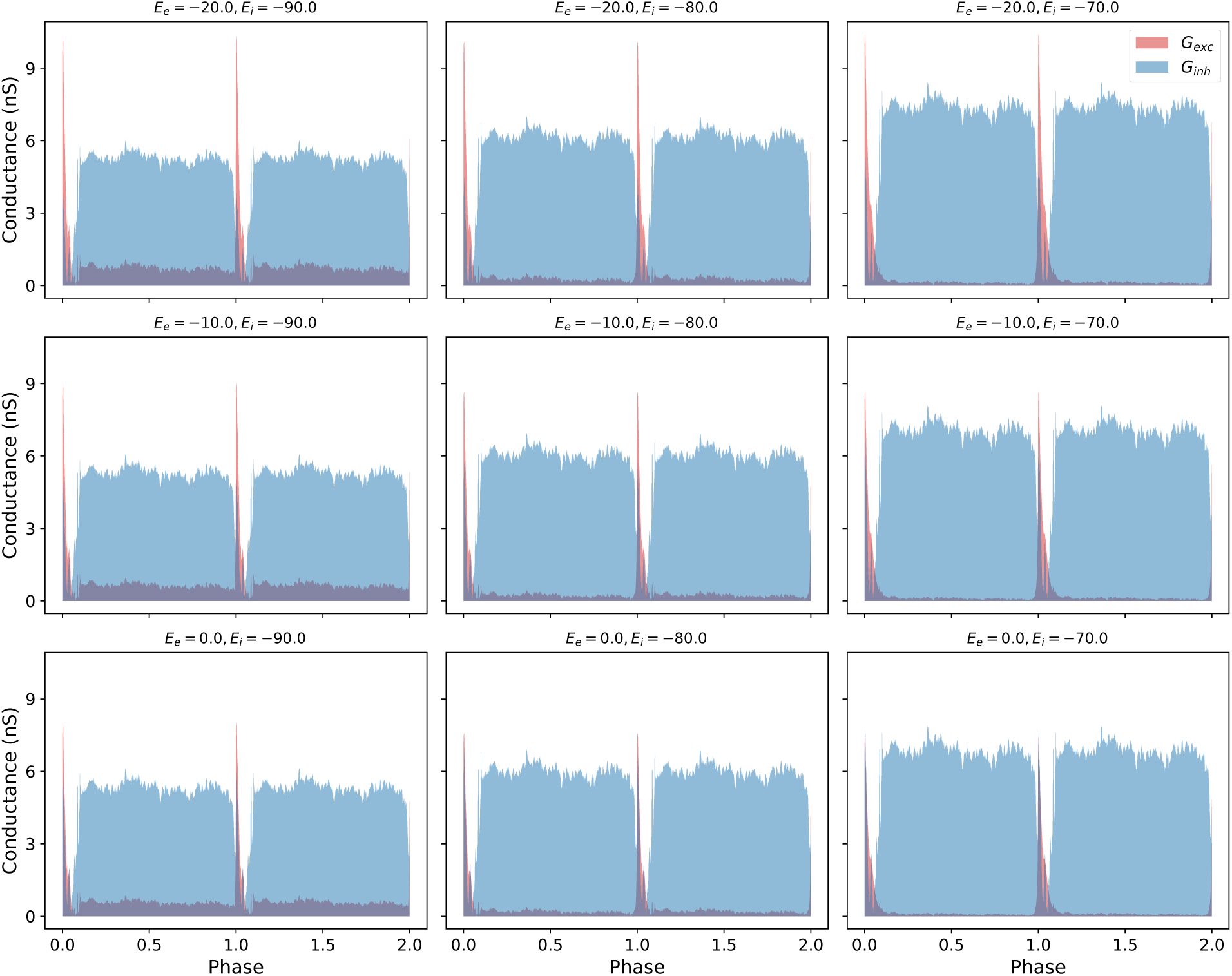
**Sensitivity of reconstructed conductances to variations in reversal potentials (*E_e_* and *E_i_*)**. Reconstructed excitatory (*G_exc_*, red) and inhibitory (*G_inh_*, blue) conductances for a representative VgluT2 inspiratory neuron were recalculated over two normalized respiratory cycles while varying the assumed excitatory and inhibitory reversal potentials. Rows show *E_e_* values of -20, -10, and 0 mV; columns show *E_i_* values of -90, -80, and -70 mV, spanning ±10 mV around the default values used for this sensitivity test. Automatic estimation of *E_i_* was disabled for this diagnostic analysis so that each panel reflects the specified reversal-potential pair. The parameter sweep primarily rescales conductance amplitudes while preserving the temporal organization of the inferred synaptic inputs, including phasic inspiratory excitation and tonic expiratory inhibition.

The amplitude of the reconstructed inhibitory conductance (*G_inh_*) scales with the assumed value of *E_i_*. As *E_i_* is shifted from -90.0 mV to -70.0 mV (moving left to right across the grid), the absolute magnitude of the calculated *G_inh_* increases. This scaling occurs as a function of the reduced driving force (*V_m_ – E_i_*) required to produce the observed current-voltage relationships. In all tested configurations, the temporal structure of the inhibition remains identical, showing a high-amplitude tonic component throughout the expiratory phase and a slight dip at the onset of the inspiratory burst.

Variations in the excitatory reversal potential (*E_e_*) result in proportional scaling of the excitatory peaks. Moving from the top row (*E_e_* = -20.0 mV) to the bottom row (*E_e_* = 0.0 mV), the peak amplitude of *G_exc_* during the inspiratory phase decreases. The timing of the excitatory burst relative to the respiratory cycle phase is preserved across all combinations of reversal potentials. The “synaptic fingerprint” of the neuron, characterized by the absence of significant excitation during the expiratory interval and the presence of a sharp inspiratory excitatory spike, is maintained in every panel of the sensitivity analysis.

The relationship between the conductances across the respiratory cycle is consistent regardless of the specific parameters chosen. The tonic nature of the expiratory inhibition persists across the entire tested parameter space. Furthermore, the relative dominance of the inhibitory conductance during the inter-burst interval compared to the excitatory conductance is a constant feature across all nine panels. The temporal organization of the synaptic inputs, including the phase-locking of the excitatory peaks and the duration of the expiratory inhibition, shows no variation across the grid of reversal potentials.

### Linearity of I-V Regressions and Cross-Mode Consistency

The validity of the conductance reconstruction relies on the linearity of the relationship between membrane current and potential over the analyzed range. Supplemental Figure 2 displays the current-voltage (I-V) relationships for all representative cell types and recording modes discussed in the primary text. Data points from two distinct phases—peak inspiration (*ϕ* ≈ 0, red) and mid-expiration (*ϕ* ≈ 0.5, blue)—are fitted with linear regressions to estimate the total synaptic conductance and the current intercept. Across all neurons, including VgluT2 and VGAT populations, the observed current values follow a linear relationship with the membrane potential, supporting the steady-state assumption of the current balance equation used for the inference.

The analysis of these I-V plots reveals high consistency between current-clamp (left panels) and voltage-clamp (right panels) recording configurations. In both modes, the slopes of the regressions undergo phase-dependent shifts that correspond to the dynamic changes in total membrane conductance. For inspiratory neurons, the separation between the red and blue regression lines illustrates the recruitment of phasic excitatory drive and the modulation of inhibitory conductance. For expiratory neurons, the steepness of the slopes during inspiration (red) reflects the strong phasic inhibition received by these cells. The high degree of linearity observed in the scatter points for both recording modes indicates that the inferred synaptic conductances are not significantly distorted by non-linear intrinsic currents within the sampled voltage ranges.

**Supplemental Figure 2.**
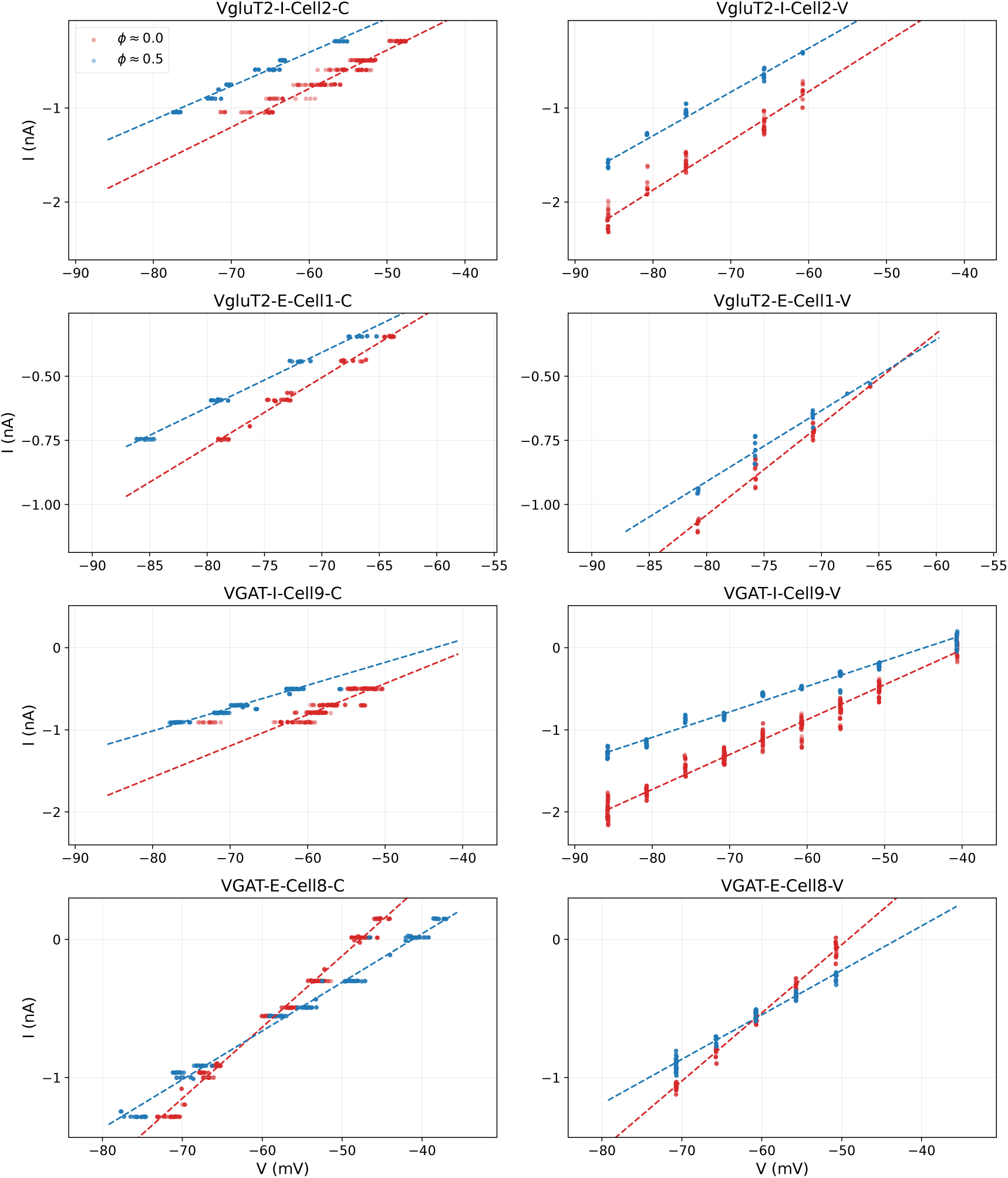
Linearity of phase-binned current–voltage relationships across populations and recording modes. Each row shows *I*–*V* data for a representative neuron of one of the four populations (top to bottom: VgluT2-I, VgluT2-E, VGAT-I, VGAT-E; cell identifier indicated above each panel). Left column: current-clamp recording. Right column: voltage-clamp recording from the same neuron. Scatter points are individual current–voltage samples drawn from two narrow phase bins of the network respiratory cycle—peak inspiration (*ϕ* ≈ 0, red) and mid-expiration (*ϕ* ≈ 0.5, blue)—and dashed lines are the corresponding robust linear fits used to estimate the total membrane conductance and current intercept (cf. Figure 2). Within the sampled voltage range, the data are approximately linear in all panels, supporting the steady-state linear current balance assumption used in the conductance decomposition. Phase-dependent shifts in slope and intercept between the red and blue regressions reflect the recruitment of phasic excitatory and inhibitory conductances over the cycle.

**Supplemental Figure 3.**
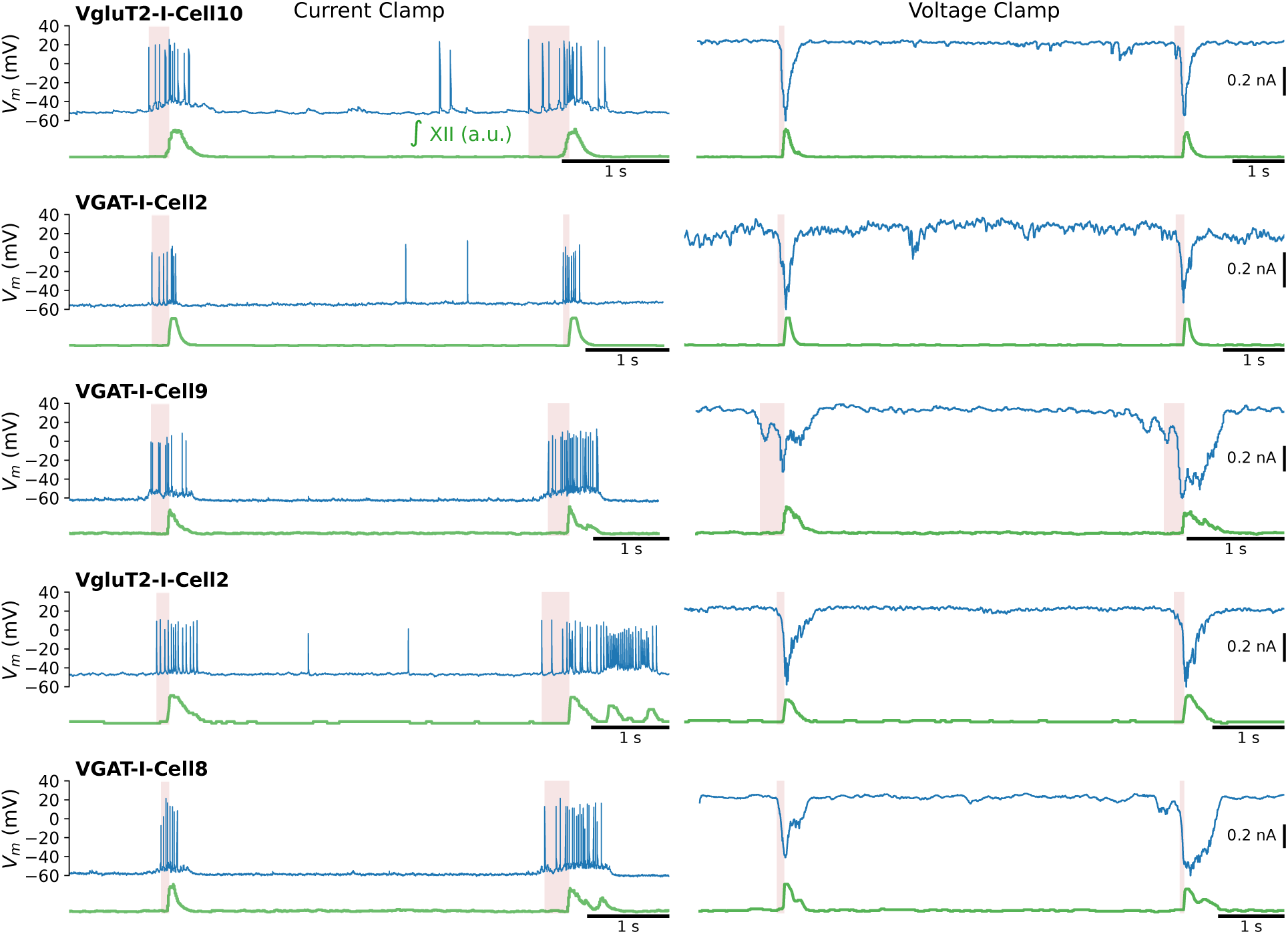
**Pre-Inspiratory Activity**. High-resolution comparison of pre-inspiratory action potential firing and underlying synaptic drive. Left column: Current-clamp episodes show pre-inspiratory spiking (blue) preceding each network burst in the reference signal (green). Right column: Corresponding median-filtered voltage-clamp holding-current traces from the same cells show the pre-inspiratory inward current (blue) preceding the reference bursts (green). Each panel contains two adjacent reference bursts; pink shading marks the pre-inspiratory lead interval for each burst. Horizontal scale bars indicate 1 s, and Voltage Clamp panels include vertical current scale bars in native units.

**Supplemental Figure 4.**
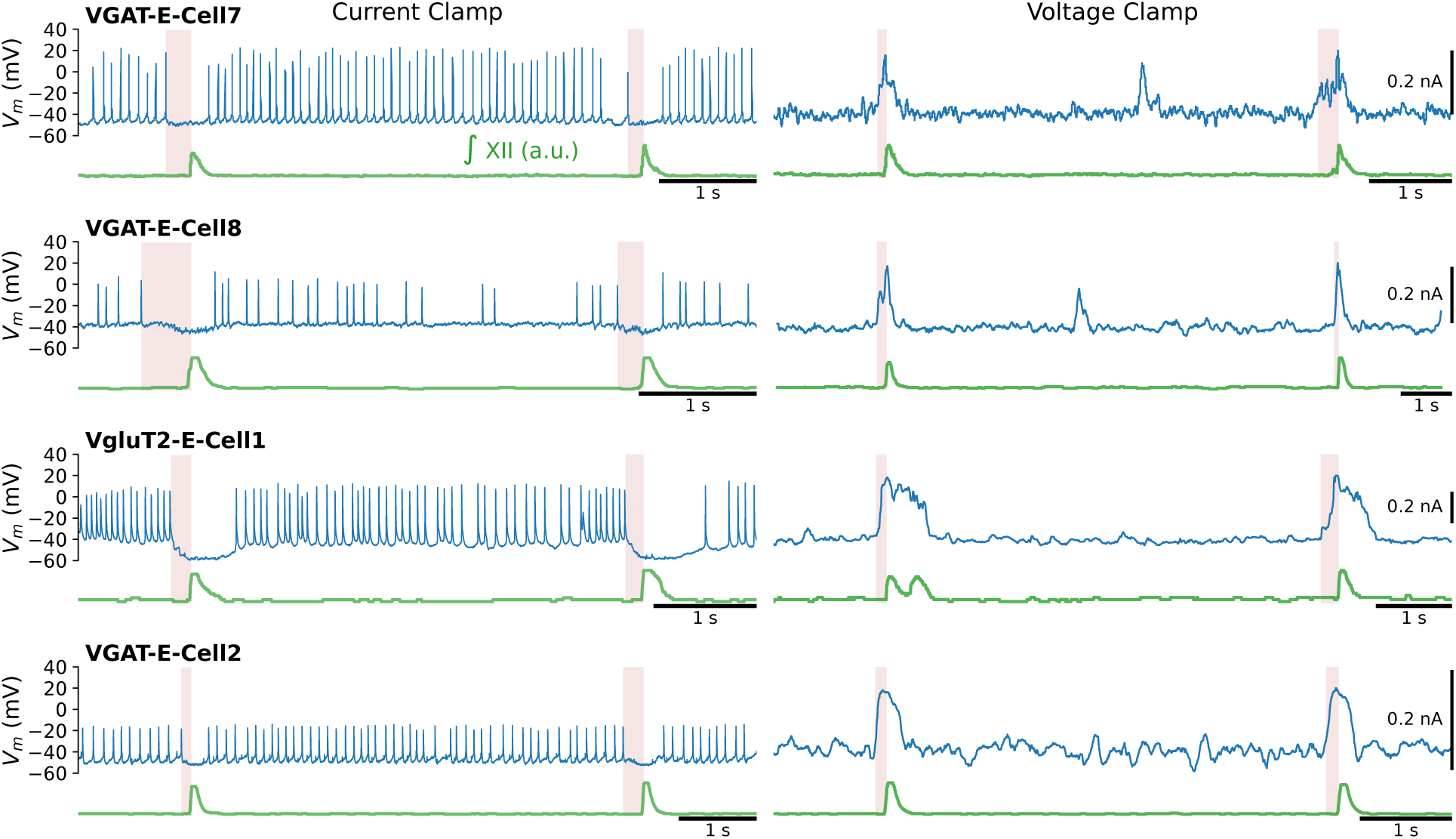
Pre-Inspiratory Inhibition of Expiratory Cells. Examples of expiratory cells that stop firing before inspiratory onset and matched voltage-clamp recordings from the same cells showing a pre-inspiratory outward current. Left column: Current-clamp episodes show expiratory spiking (blue) terminating before each inspiratory burst in the reference signal (green). Right column: Corresponding median-filtered voltage-clamp holding-current traces show an outward synaptic current (blue) preceding the same inspiratory reference bursts (green). Each panel contains two adjacent reference bursts; pink shading marks the pre-inspiratory silent interval in current clamp and the pre-inspiratory outward-current interval in voltage clamp. Horizontal scale bars indicate 1 s, and voltage-clamp panels include vertical current scale bars in native units.

### Pre-Inspiratory Dynamics and Phase Transitions

These supplemental panels provide representative examples of the pre-I dynamics discussed in the main text.

Pink intervals in Supplemental Figures 3–4 mark the interval between a cell-specific pre-I event and the onset of the next population inspiratory burst. The right edge of each interval is the onset of the next network inspiratory burst, detected from the smoothed integrated XII reference and aligned to its sharp rising phase. In Supplemental Figure 3, the left edge marks the beginning of pre-I spiking in current clamp or the first sustained inward deflection in the filtered voltage-clamp holding current. In Supplemental Figure 4, the left edge marks the end of expiratory spiking in current clamp or the first sustained outward deflection in the filtered voltage-clamp holding current.

## Notes

https://ymolkov.github.io/synaptic-inputs-slices/

https://github.com/ymolkov/synaptic-inputs-slices

### Competing Interest Statement

The authors have declared no competing interest.

